# Molecular dissection of the soluble photosynthetic antenna from a cryptophyte alga

**DOI:** 10.1101/2023.08.08.552318

**Authors:** Harry W. Rathbone, Alistair J. Laos, Katharine A. Michie, Hasti Iranmanesh, Joanna Biazik, Sophia Goodchild, Pall Thordarson, Beverley R. Green, Paul M. G. Curmi

## Abstract

Cryptophyte algae have a unique phycobiliprotein light-harvesting antenna that fills a spectral gap in chlorophyll absorption, however, it is unclear how it transfers energy efficiently to photosystems. We show that the cryptophyte *Hemiselmis andersenii* expresses an energetically complex antenna comprising three distinct spectrotypes of phycobiliprotein with different quaternary structures arising from a diverse α subunit family. The bulk of the antenna consists of *open* quaternary form phycobiliproteins acting as primary photon acceptors, supplemented by novel *open-braced* forms. The final components are *closed* forms with a long wavelength spectral feature due to substitution of a single chromophore. We propose that the macromolecular organization of the cryptophyte antennas consists of bulk *open* and *open-braced* forms that transfer excitations to photosystems via this bridging *closed* form phycobiliprotein.

**One-Sentence Summary:** Algae generate a rainbow of antenna proteins by combining a conserved subunit with different members of a multigene family.

## Main Text

Cryptophytes are single-celled algae that obtained their chloroplasts by secondary endosymbiosis of a red alga (*1-3*). The ancestral red algal light harvesting antenna, the phycobilisome (PBS), is a 1.2-20 MDa complex attached to the stromal side of the thylakoid membranes which creates an energetic funnel from the outer reaches of the complex down to the integral membrane photosystems (*4, 5*). In contrast, the cryptophyte antenna is comprised of repurposed phycobilisome components (*3*) densely-packed between the photosynthetic membranes in the thylakoid lumen.

Each cryptophyte phycobiliprotein (PBP) is a (hetero)dimer of two αβ protomers where each protomer contains a unique α subunit and a common β subunit, where the latter has descended virtually unchanged from the red algal PBS phycoerythrin (PE) β protein (*6*). Crystal structures show that the cryptophyte PBPs take on two distinct quaternary structures: the *closed* form (*7, 8*) and the *open* form (*9*). The latter form results from a single amino acid insertion in the α subunit which alters the packing between αβ protomers, creating a large solvent filled cavity (*9*). Sequence data shows that *open* forms exist only in the *Hemiselmis* genus (*10*).

Since the discovery of the cryptophyte light harvesting system (*11-13*), it has been widely believed that the soluble antenna is comprised largely of copies of a single PBP per organism with a specific absorption maximum between 545 and 645 nm giving each PBP its name (e.g. PE545) (*1, 14-16*). This is despite evidence to the contrary coming from absorption spectroscopy and isoelectric focusing (*11-13, 17-22*) as well as evidence of multiple nuclear α subunit genes from the genome sequence of *Guillardia theta* plus transcriptomic data (*17, 23*). In fact, all 20 α subunit genes in *G. theta* are expressed in culture (*24*). Finally, for many cryptophytes there appears to be an impassable spectral gap between the fluorescence emission of the major PBPs and the 670-710 nm absorption maxima of the membrane-bound photosystems.

In this paper we demonstrate that the cryptophyte *Hemiselmis andersenii* has an energetically complex antenna with multiple protein components possessing different spectral and structural characteristics, along with a new quaternary structure. Using energetic considerations, we define specific roles for each of these proteins in order to generate a functional antenna and formulate a simple model for the organization of this antenna *in vivo*. Finally, we show that this model appears to be general, harkening back to previous studies on other cryptophytes with unexplained spectral features.

Analysis of the transcriptomes of four strains of *H. andersenii* yielded 22 unique cryptophyte α subunit sequences (Fig. 1a; Table S1; Materials & Methods) (*17, 18*). The final mature sequence set consists of eight *closed* forms (*Ha*α^C^_1_ to *Ha*α^C^_8_, Fig. 1a) and eleven *open* forms (*Ha*α^O^_1_ to *Ha*α^O^_11_) (Fig. 1a). Three additional *open* forms contain a seven-residue insertion between the second stand of the β ribbon (*S2*) and the α helix (*H1*) and these three sequences are here termed ‘*open-braced*’ (*Ha*α^OB^_1_to *Ha*α^OB^_3_; Fig. 1a). Two *open* forms, *Ha*α^O^_3_ and *Ha*α^O^_4_, had smaller incursions in this region (Supplementary Text D). The eight *closed* forms fit largely into distinct αL (‘long’; *Ha*α^C^_1_ and *Ha*α^C^_5_) and αS (‘short’; *Ha*α^C^_2_, *Ha*α^C^_3_, *Ha*α^C^_4_ and *Ha*α^C^_6_) categories (*10*). Thus, after the emergence of *open* forms in the *Hemiselmis* genus (*9*), closed forms still remain suggesting a function reason.

**Figure 1.**
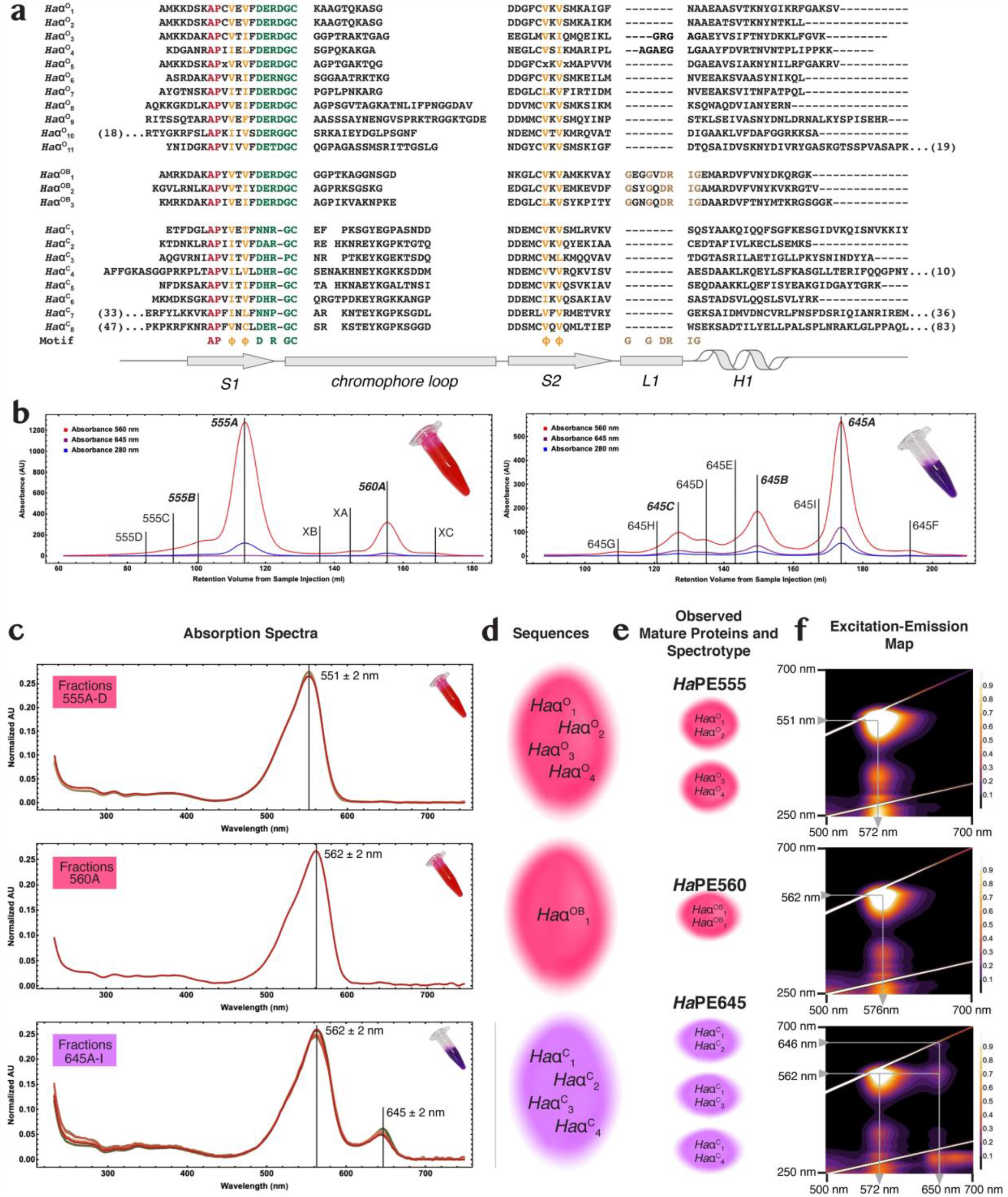
Hemiselmis andersenii soluble antenna is composed of multiple PBPs that fall into three spectrotypes. **a**. Alignment of all *Hemiselmis andersenii* α subunit transcripts, clustered by *open, open-braced* and *closed* state predicted quaternary structures. The structural motif is at the bottom. Color coding: red – identical; orange – conserved property; green -chromophore site; and brown – *open-braced* form *L1* motif. Numbers in parentheses indicate additional N- and C-terminal residues. **b**. Cation exchange chromatograms show the zoo of light harvesting proteins as peaks in chromatograms. Two fractions: ‘pink’ and ‘purple’ (see images inset in chromatograms) were first separated by anion exchange chromatography. The left chromatogram is for the ‘pink’ fraction while the right chromatogram is for the ‘purple’ fraction. The largest peaks are shown in bold with all peaks labelled by their spectrotype. The chromatograms show absorbance at three visible wavelengths: 560nm red; 645nm purple; and 280nm blue. **c-f**: Diagram showing the flow of characterization for peaks isolated in **b** from: **c**. absorption spectra of peaks to **d**. identification of α subunit components via fragment LC-MS/MS to **e**. identification of purified mature proteins via intact mass spectrometry to **f**. characterization of spectrotypes by fluorescence emission-excitation maps. **c**. Overlays of visible absorption spectra for each chromatographically separated species belonging to the three principal spectrotypes. The *Ha*PE555 (top) and *Ha*PE645 (bottom) panels contain an overlay of multiple spectra from chromatographically distinct peaks, while the *Ha*PE560 (center) panel contains a single spectrum. **d**. Transcript sequences that have been identified via fragment LC-MS/MS to correspond to peaks in the chromatograms sorted by spectrotype. **e**. These sequences are assembled into mature proteins where the component α subunits are paired based on the highest scoring observations by intact mass spectrometry. **f**. Excitation-emission maps for the most abundant protein from each of the three spectrotypes. The arrows map the energy transfer pathways through each protein. The vertical axis is the excitation wavelength and the horizontal axis is the emission wavelength. The highest peaks have been truncated so as to highlight the smaller features in the heat maps.

The soluble *H. andersenii* PBPs were extracted and first partitioned into two distinct colored fractions, one pink, the other purple, by anion exchange chromatography (Fig. 1b left and right, respectively; Fig. S1; Materials & Methods). Each of these two fractions was further separated into a number of discrete protein fractions by cation exchange chromatography, that are labelled according to their distinguishing wavelength maximum and a letter indicating its relative abundance (Fig. 1b).

Linear absorption spectra show that each isolate fell into one of three distinct categories (Fig. 1c; Supplementary Text A). The major spectrotype, *Ha*PE555, has an absorption peak at 551 ± 2 nm (Fig. 1c top row); the second, *Ha*PE560, has an absorption peak at 562 ± 2 nm (Fig. 1c middle row); and the last, *Ha*PE645, has a peak at 562 ± 2 nm with a smaller secondary peak at 645 ± 2 nm (Fig. 1c bottom row).

Peptide identification by fragment LC-MS/MS and intact mass spectrometry were combined to match sequence data with purified protein fractions. Nine α subunit sequences were identified with high confidence by their MASCOT score (Fig. 1d; Materials & Methods). Using intact mass spectrometry, the most probable pairings to produce mature α_1_β.α_2_β PBPs were: *Ha*α^O^_1_— *Ha*α^O^_2_ (fraction 555A), *Ha*α^OB^_1_—*Ha*α^OB^_1_(560A), *Ha*α^C^_1_—*Ha*α^C^_2_ (645A and 645B), *Ha*α^C^_1_— *Ha*α^C^_3_ (645B and 645C), and possibly *Ha*α^O^_3_— *Ha*α^O^_4_ (XA) and *Ha*α^C^_1_—*Ha*α^C^_4_ (645C) (Fig. 1e; Materials & Methods).

Abundances for each light harvesting protein were estimated by comparing peak areas in chromatograms (Materials & Methods). Each spectrotype appears to have one clear majority fraction. The largest peak of *Ha*PE555 (555A) is by far the most abundant PBP constituting approximately 63% of the overall protein complement (with other *Ha*PE555 spectrotype peaks contributing 8%). The *Ha*PE560 peak constitutes 14% of the protein complement and the largest peak of *Ha*PE645 constitutes 8% (with other *Ha*PE645 spectrotype proteins contributing 7%; Table S2). This suggests that the spectrotypes are approximately in the ratio of 5:1:1 (*Ha*PE555:*Ha*PE560:*Ha*PE645). These data show that there is an overwhelming majority of a single PBP in the antenna of *H. andersenii*, that of spectrotype *Ha*PE555, explaining why the one-PBP-per-organism convention has persisted.

Fluorescence excitation-emission maps were produced for each spectrotype to provide clues to the overall energetic architecture of the antenna (Fig. 1f; Materials & Methods). These maps show that *Ha*PE560 and *Ha*PE555 spectrotypes have tightly peaked spectra with similar excitation and emission wavelength ranges (Fig. 1f; *Ha*PE555: excitation peak 551 ± 2nm, emission peak 572 ± 1nm; *Ha*PE560: excitation peak 562 ± 2nm, emission peak 576 ± 1nm). *Ha*PE645 has many of the same features as the other two spectrotypes (excitation peak 562 ± 2nm and major emission peak 572 ± 1nm), however, it has an extra peak absorbing at 646 ± 2nm and emitting at 650 ± 1nm plus a cross-peak connecting the two wavelength regions (excitation at 562 ± 2nm, emission at 650 ± 1nm). The excitation-emission map for *Ha*PE645 suggests that this protein acts as an energy adaptor with excitation transfer from higher energy (562nm) to lower energy (650nm) chromophores within the protein.

Four new structures of *Ha*PE555 were determined (Fig. S2a; Table S3; resolutions: 1.57Å, 1.67Å, 1.73Å and 1.95Å). Three of these (PDB: 8EL3, 8EL4, 8EL5) were derived from peak 555A (Fig. 1b left) and the remaining one (PDB: 8EL6) was derived from peak 555B (Fig. 1b). They all have the *open* form quaternary structure with two distinct, but near identical α subunits (Fig. S2a bottom). Each protein in the new structures matches that of 4LMX to RMSD 0.46 ± 0.17 Å over 2937 ± 124 atoms, however, they differ in their packing and a small alteration in secondary structure in the structure from chromatogram peak 555B (8EL6), (Supplementary Text B; Fig. S3).

All five crystal forms of protein *Ha*PE555 are constructed from continuous filaments of PBPs, which are organized into two-dimensional rafts (Fig. S2b; Supplementary Text C). The filaments are generated via a knob and hole packing but differ in the distances between adjacent molecules along the filament (having either tight or loose interfaces; Fig. S2b; Table S4). This packing brings the chromophores of neighboring PBPs closer together increasing the possibility of energy transfer (Fig. S2c-e). This filament arrangement appears to be unique to *Ha*PE555, although filaments have been observed for the *closed* forms: *Cp*PE566 from *Cryptomonas pyrenoidifera* (*10*); and *Ps*PE545 from *Proteomonas sulcata* (*25*).

The crystal structure of *Ha*PE560 was determined at 1.45Å resolution (PDB 7SSF, Fig. 2a; Table S3). It shows a symmetric (αβ)_2_ dimer (Fig. 1e middle; Fig. 2a) that resembles the *open* form structure of *Ha*PE555 with which it shares the same chromophore arrangement despite the spectral differences (Table S5; Fig. 1c). However, the *Ha*PE560 structure shows several critical differences. The key structural difference between *Ha*PE560 and *Ha*PE555 is that the *Ha*PE560 α subunit contains an inserted loop of seven residues (*L1*) between β-strand *S2* and α-helix *H1* (Fig. 1a). This *L1* loop partially closes the entrance to the central solvent filled cavity of *Ha*PE560 when compared to *open* form PBPs such as *Ha*PE555 (Fig. 2a-c; Fig. S4a). In the *open* form, this cavity is created by the insertion of an Asp residue just prior to the chromophore binding Cys residue (*9*) where this change in quaternary structure reduces the buried solvent accessible surface area between two αβ protomers to 1,084 ± 63 Å^2^ for *Ha*PE555 compared to 2,191 Å^2^ for the *closed* form of *Chroomonas Cs*PC645. In *Ha*PE560, the insertion of the *L1* loop redresses this loss of interaction by increasing the buried surface area between two αβ protomers to 1,452 ± 2 Å^2^. Thus, the evolution of the *L1* loop strengthens and stabilizes the interface between adjacent α subunits connecting the two αβ protomers, thereby consolidating the *open* form quaternary structure. Given this, we call the new quaternary structure of *Ha*PE560 the ‘*open-braced*’ form.

**Figure 2.**
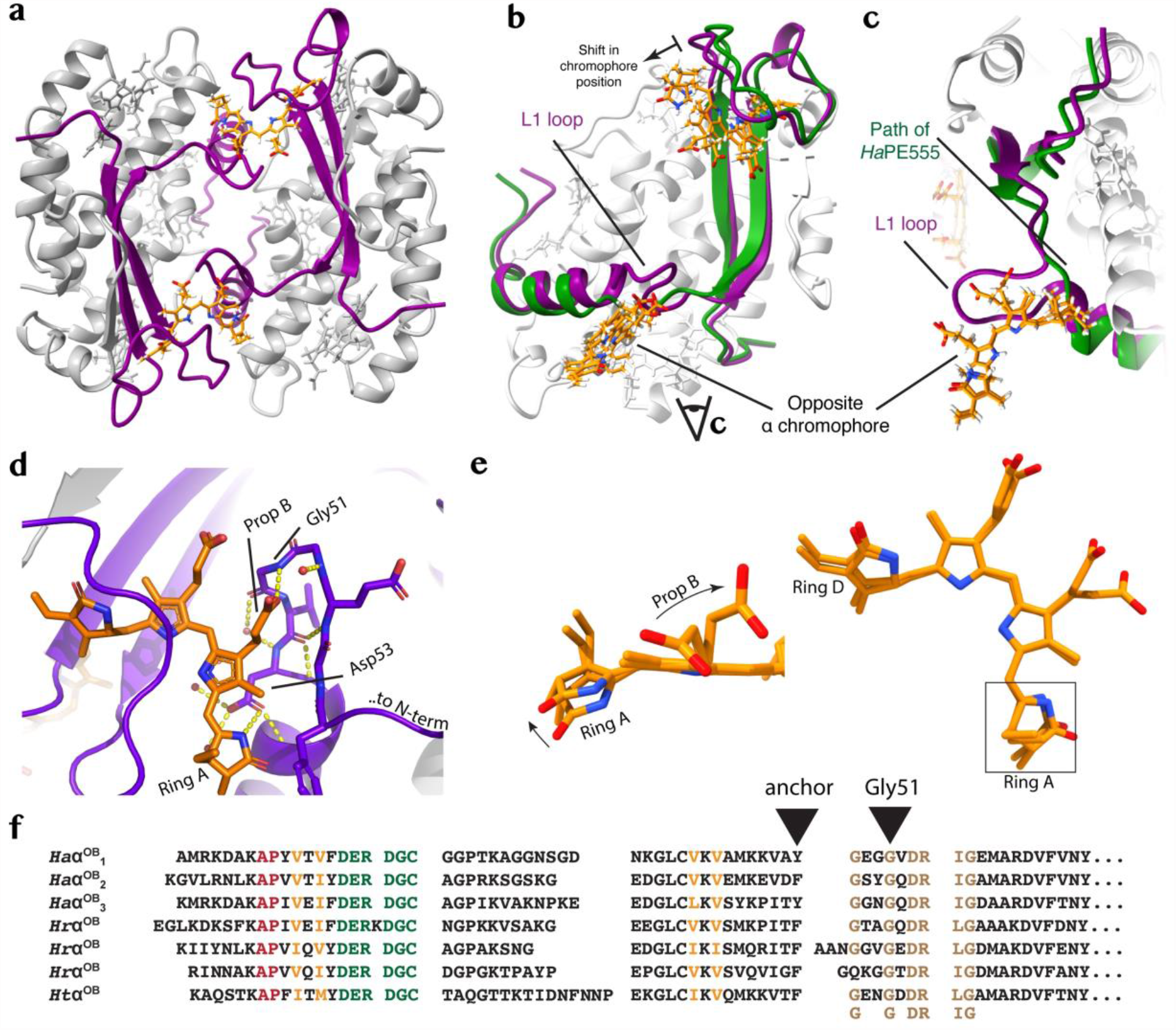
The crystal structure HaPE560. **a**. Cartoon diagram showing the crystal structure of the *open-braced* quaternary form *Ha*PE560 (α subunits purple, β subunits gray). **b**. Comparison of the *open-braced Ha*PE560 α subunit structure with that of the *open* form *Ha*PE555. The *open-braced* α subunit is shown in purple cartoon while the *open* form α subunit is in green. The *L1* loop can be seen in the center of the panel. **c**. Inset comparing the *L1* protrusion (marked) and the path of the *Ha*PE555 (green) as viewed by the eye marker in **b**. The chromophores of both proteins are shown in gold in **b** and **c**, with the chromophore of *Ha*PE560 shifted away from the binding pocket. **d**. The interaction between the *L1* loop and the neighboring α chromophore. The seven residue *L1* loop interacts with the α19 chromophore of the opposite α subunit in two key places: Asp53, which ligates the nitrogen of pyrrole ring A and Gly51, where the propionate from pyrrole ring B forms a hydrogen bond to the backbone nitrogen, stabilizing the β hairpin structure. **e**. The interaction between the *L1* loop and the α19 chromophore from the opposite α subunit leads to: a rotation of pyrrole ring A relative to ring B, when compared to *open* form structures; and an anchoring of the propionate side chain of pyrrole ring B, decreasing flexibility (and thus excitation decay routes). Right panel shows the complete α19 chromophore in the same orientation as **c**, while the left panel highlights the changes in chromophore geometry. **e**. An alignment of *open-braced* form α subunit transcriptome sequences from: *Ha* – *H. andersenii*; *Hr* – *H. rufescens*; and *Ht* – *H. tepida*. Color codes are asp per Fig. 1a.

The *L1* loop insertion alters the conformation and geometric arrangement of the α chromophore in the *open-braced* quaternary structure. The *L1* loop forms a twisted β hairpin and scaffolds the opposite α chromophore (Fig. 2abc; Fig. S5a). The propionate group emanating from pyrrole ring B forms a hydrogen bond to the backbone nitrogen of Gly51 forming the tip of the *L1* hairpin (tight-turn), stabilizing the hairpin structure (Fig. 2d). The side chain of Asp53 on the descending strand of the *L1* β hairpin contacts the opposite α chromophore and forms a hydrogen bond to the nitrogen of pyrrole ring A (Fig. 2d). This interaction with Asp53 pulls pyrrole ring A away from the usual *open* form conformation and rotates pyrrole ring A (Fig. 2e; Fig. S6a-c). It is therefore likely responsible for the ∼10 nm wavelength shift of *Ha*PE560 absorption compared to *Ha*PE555 (which has identical chromophore placements) (Fig. 2b-e; Fig. S6a-c).

Examination of transcriptome data for *Hemiselmis tepida* and *Hemiselmis rufescens* shows that each of these organisms contains α subunit sequences that show a homologous *L1* loop insertion that is a signature for the *open-braced* form (Fig. 2f; Supplementary Text D). The transcriptomes for these organisms also contain multiple *open* form α subunit sequences, out numbering the *open-braced* form (Fig. S7; *open*:*open-braced* transcripts – 11:3, 8:6 and 8:1 for *H. andersenii, H. refescens* and *H. tepida*, respectively). Thus, the more stable *open-braced* form has not displaced the *open* form, suggesting there is a functional reason for maintaining these two forms.

The structure of the *closed* form *Ha*PE645 provides an explanation for its unusual spectral properties. The crystal structure of the major *Ha*PE645 species (Fig. 1b) was determined at 1.49Å resolution (PDB 7SUT; Fig. 3a; Table S3) and shows a typical *closed* form PBP (*7-9*) with αL and αS sequences corresponding to *Ha*α^C^_1_ and *Ha*α^C^_2_, respectively (Fig. 1a). Our electron density maps clearly demonstrate that the chromophore attached to Cys82 on the β subunit in the αSβ protomer is unambiguously PCB (which causes the secondary absorption peak for this protein at 645 nm; Fig. 3b), while the chromophore attached to Cys82 on the β subunit in the αLβ protomer is unambiguously PEB (Fig. 3c). Such an asymmetry has not been observed previously for cryptophyte β subunits within a single PBP and poses an interesting question – how can two identical β-subunits within the same organism acquire different chromophores at position 82?

**Figure 3.**
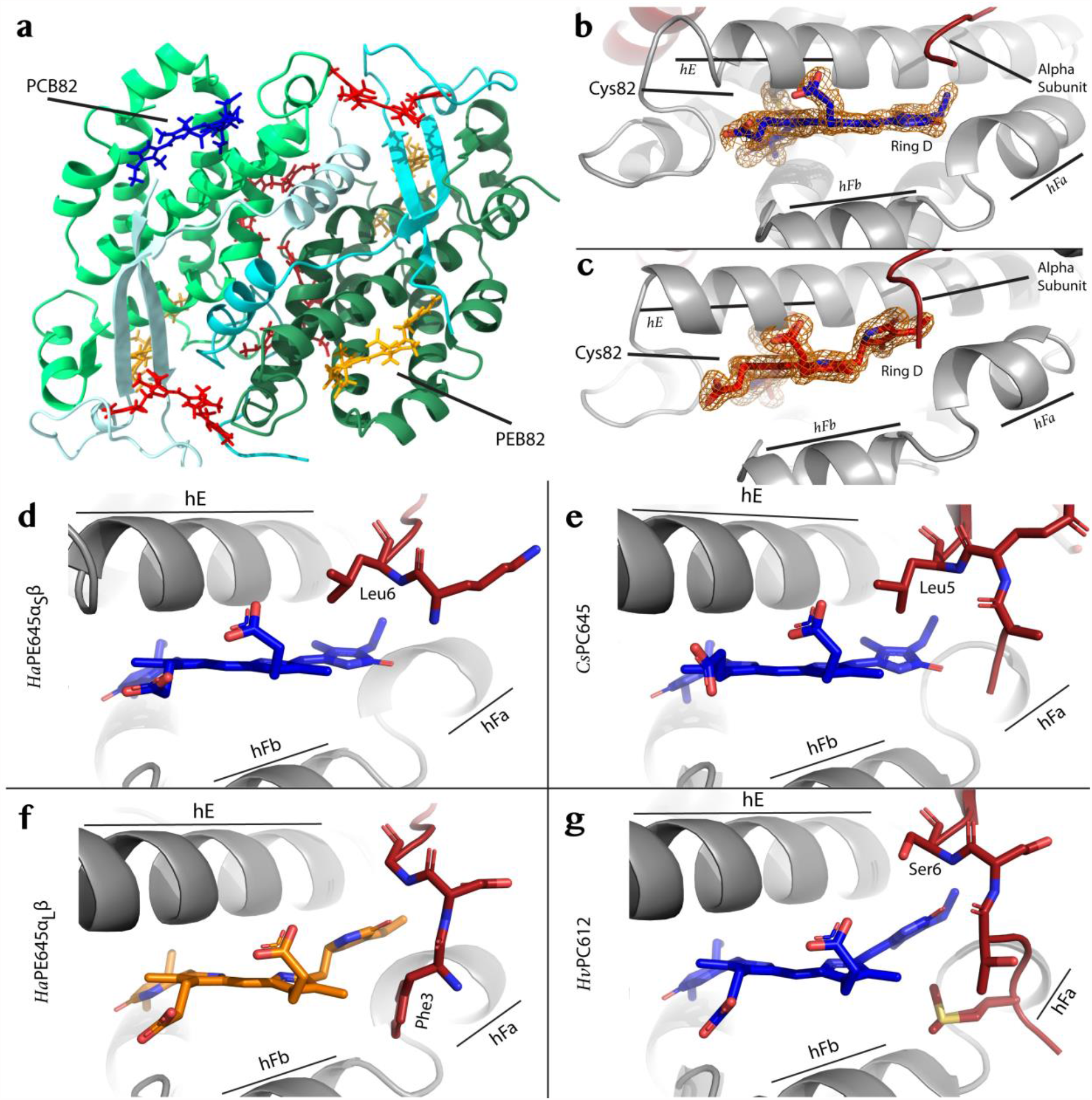
The crystal structure of HaPE645. **a**. Cartoon representation of the *closed* form *Ha*PE645 structure (αL cyan; αS pale cyan; corresponding β subunits forest and lime green, respectively). **b**. The electron density (orange mesh) for the β82 chromophore attached to the αSβ protomer clearly shows that it is PCB, where pyrrole ring D is part of the conjugated structure. **c**. In contrast, the electron density (orange mesh) for the β82 chromophore attached to the αLβ protomer clearly shows that it is PEB, where pyrrole ring D is separated by two sp3 carbon atoms from the conjugated ring structure. **d**. Shows the interactions between the N-terminus of the αS subunit and the PCB β82 chromophore in *Ha*PE645, which are similar to those seen in **e**. for the PCB β82 chromophore in *Cs*PC645. **f**. Shows the interactions between the N-terminus of the αL subunit and the PEB β82 chromophore in *Ha*PE645, which are similar to those seen in **g**. for the PCB β82 chromophore in *Hv*PC612.

The N-terminal region of the α subunit appears to dictate the chemical nature of the chromophore attached to Cys82 on the associated β subunit. In each αβ protomer, the N-terminal tail of the respective α subunit interacts with pyrrole ring D of the β82 chromophore. For the αSβ protomer, the side chain of Leu6 packs against the face of pyrrole ring D of PCB β82 (Fig. 3d). This pocket is specific for PCB and incompatible with a PEB chromophore due to the steric clash that would be created by Leu6 (Fig. 3d compared to Fig. 3f). There is a similarity between *Ha*PE645 and *Cs*PC645 when looking at the residues surrounding the β82 PCB chromophore (compare Fig. 3d to 3e, respectively). The conformation of PCB β82 is nearly identical when comparing the αSβ protomer of *Ha*PE645 to both β82 sites in *Cs*PC645 (*9*). At all three sites, leucine makes an equivalent interaction with PCB β82 pyrrole ring D (Fig. 3de). In other cryptophyte-PC forms (*Hv*PC612, *Cs*PC630 and *Hp*PC577), this residue is replaced by a glutamic acid or serine which induces a rotation of pyrrole ring D of the chromophore, tuning the absorption away from 645 nm (*9, 10*) (Fig. 3g; Fig. S6de; Supplementary Text E).

How can two identical β-subunits acquire different chromophores at position 82? Lyase enzymes mediate the stereospecific ligation of the correct chromophore to a specific cysteine (*26*). The cryptophyte *G. theta* has been shown to possess members of each family of bilin lyase including an S-type lyase which is specific for attaching a chromophore to Cys82 (*27*). The S-type lyases transfer chromophores to a folded apo-PBP (*28*). Our data indicate that the substrate for the Cys82 lyase is the folded apo αβ protomer, as the N-terminus of the α subunit confers chromophore specificity. It remains to be discovered whether *H. andersenii* possesses two S-type lyases, one for PCB and one for PEB, given the asymmetry in *Ha*PE645 (Supplementary Text F).

A recent study of photoacclimation in two cryptophyte species has shown that the absorbance spectra of the PBPs are altered when cells are photoacclimated via growth under restrictive light conditions using spectral filters (*25*). This study concluded that the spectral changes must result from differential chromophorylation of the PBPs which is consistent with the existence of a pool of lyases capable of modifying the attached chromophores as observed in our structure of *Ha*PE645.

Given the presence of at least three distinct PBP components in *H. andersenii*, what is the organization of this antenna? Electron micrographs (using high pressure freezing and freeze substitution) of the plastids of *H. andersenii* show clear striations of electron opaque material between the thylakoid membranes indicating densely-packed protein, as observed for other cryptophytes (*29-31*) (Fig. 4a). The width of the electron opaque material within the thylakoid lumen is estimated to be 12.7 ± 2.9 nm (Fig. 4b), which is within the 10-50 nm range of previous measurements (where the width increases with the inverse of light intensity during culturing) (*29-31*). Based on the dimensions of the proteins from our crystal structures, we estimate that only around 3-4 proteins (or up to 10-13 proteins in very low light (*29-31*)) would fit across the thylakoid lumen (Fig. 4c; Materials & Methods).

**Figure 4.**
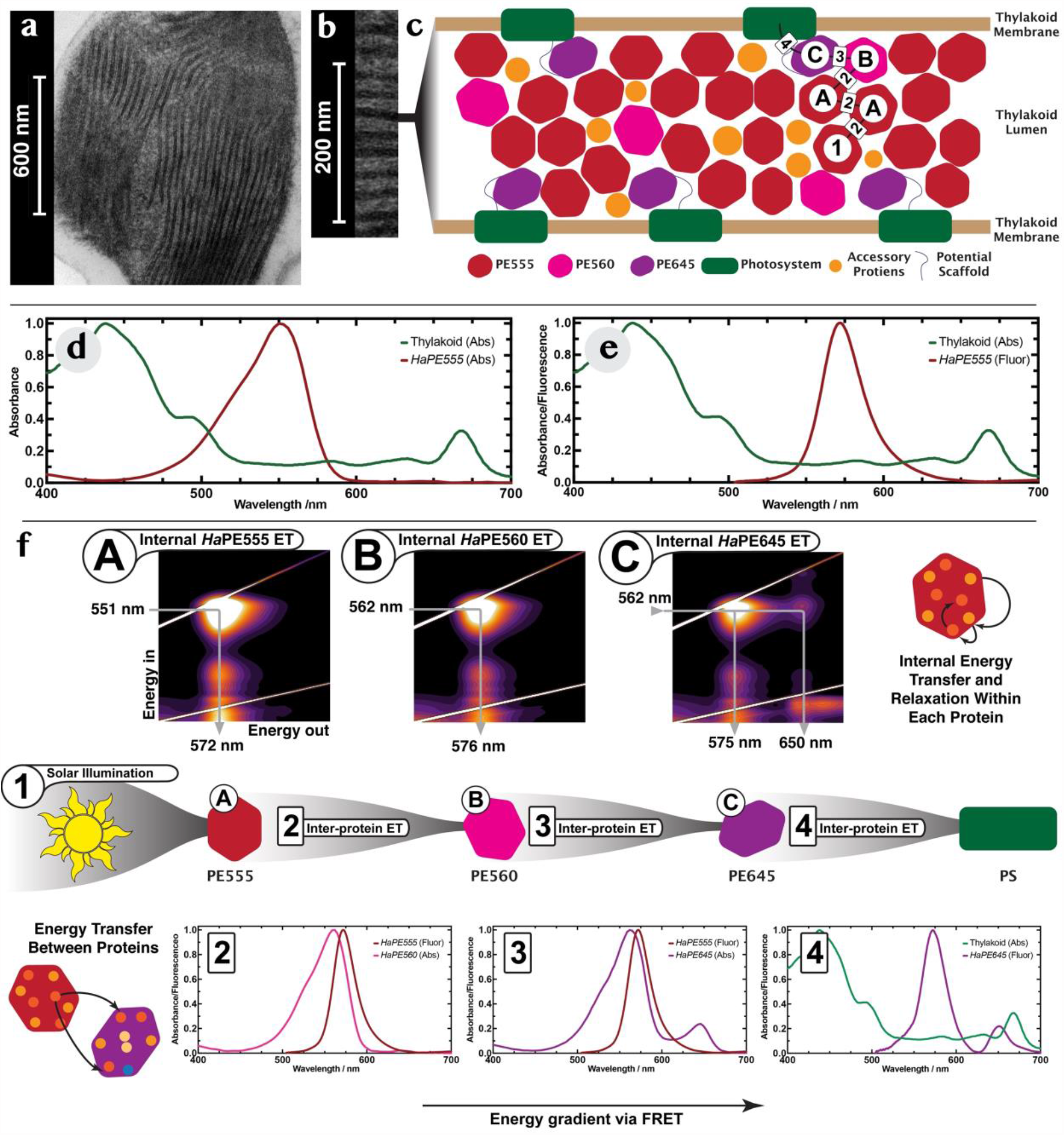
The organization of the soluble cryptophyte light harvesting antenna in the thylakoid lumen. **a**. Electron micrograph of a thin section from a whole freeze-substituted *H. andersenii* cell (overview). **b**. Close up of the chloroplast showing clear striations of dense material in the thylakoid lumen. **c**. A model of the soluble light harvesting antenna residing in the lumenal space of the thylakoid. *Ha*PE645 (purple) resides near the membrane while the lumenal space is filled with a dense phase of *Ha*PE555 and *Ha*PE560 (red and pink, respectively). A typical energy transfer path is shown as a series of black lines connecting PBPs to the integral membrane photosystems (green). The capital letters and number along this path refer to energy transfer processes as described in panel **f. d**. The absorption spectrum of the major soluble cryptophyte antenna protein *Ha*PE555 (red) plugs a spectra hole in the absorption spectrum of the photosystems and antennas in the thylakoid membrane (green). **e**. the fluorescence emission of *Ha*PE555 (red) shows minimal spectral overlap with the absorption spectrum of the thylakoid membrane system (green) where chlorophyll is the main chromophore in the integral membrane antennas and photosystems. **f**. Proposed light harvesting pathway shown as a series of coupled excitation-emission maps (**A**-**C**) demonstrating internal excitation transfer within PBPs with associated 1D spectra showing overlap between emission and excitation (**2**-**4**) for inter molecular energy transfer between proteins. This pathway corresponds to that shown in panel **c. Step 1**: sunlight is incident on *Ha*PE555 which absorbs around 551nm. Energy transfer moves the excitation within the protein (excitation-emission map **A**). **Step 2**: energy is transferred to either *Ha*PE555 or *Ha*PE560 as the emission spectra of *Ha*PE555/*Ha*PE560 has significant spectral overlap with the absorption spectra of *Ha*PE555/*Ha*PE560 (spectrum **2** shows overlap of emission of *Ha*PE555 (red) and absorption of *Ha*PE560 (pink)). The excitation moves within *Ha*PE560 (excitation-emission map **B**). **Step 3**: energy is transferred from either *Ha*PE555 or *Ha*PE560 to *Ha*PE645 as their emission spectra **3** (red, only *Ha*PE555 shown) have significant spectral overlap with the absorption spectrum of *Ha*PE645 (purple). Within *Ha*PE645, the excitation migrates to the final acceptor PCB β82 chromophore (excitation-emission map **C**). **Step 4**: spectral overlap between *Ha*PE645 emission (purple) and photosystem absorption (green) facilitates energy transfer to the integral membrane system. The location of the *Ha*PE645 adaptor that couples the soluble antenna to the integral membrane systems must be close to the membrane for efficient transport (panel **c**).

Fluorescence recovery after photobleaching (FRAP) experiments attest to the density and restricted mobility of protein in the thylakoid lumen (*32, 33*). After photobleaching, the PBP fluorescence either did not recover (no diffusion of PBP over several minutes; *Rhodomonas salina*) (*33*) or recovered very slowly giving a diffusion coefficient of (1.5 ± 0.1)×10^−9^ cm^2^/s which is significantly smaller than that of comparable proteins in other cellular compartments (*Rhodomonas* CS24) (*32*).

All the results above paint a picture of a tightly packed, energetically complex, multi-subunit protein antenna embedded in the thylakoid lumen of the cryptophyte *H. andersenii* where the predominant PBPs are *Ha*PE555, *Ha*PE560 and *Ha*PE645 in the ratio of 5:1:1. From this, how does the antenna organize itself within the thylakoid lumen for robust and efficient energy transport? We have shown that *Ha*PE555 and *Ha*PE560 form the bulk of the light harvesting antenna (roughly 6/7 of the PBPs) and capture the highest energy photons. Absorption and fluorescence derived from excitation-emission maps suggest that *Ha*PE555/560 spectrotypes can transfer excitations between each other as they have significant spectral overlap (Fig. 1f, Fig. 4f Step 2). The absorption of *Ha*PE555 and *Ha*PE560 lie in a spectral region transparent to the chlorophylls that make up the integral membrane photosystems (Fig. 4d) (*34*). Thus, *Ha*PE555 and *Ha*PE560 have evolved to capture photons that would be otherwise lost by the integral membrane photosystems: they plug a spectral hole.

However, the integral membrane photosystems of cryptophytes have very little spectral overlap with the fluorescence of *Ha*PE555/560 (Fig. 4e), thus, *Ha*PE555 and *Ha*PE560 are unlikely to transfer energy directly to the membrane systems. Hence, another element is required to bridge the energy gap between the bulk antenna (*Ha*PE555 and *Ha*PE560) and the integral membrane photosystems.

The spectral properties of *Ha*PE645, a much smaller constituent fraction of the cryptophyte PBP antenna (∼1/7 of the PBPs), fulfil the requirements for a terminal acceptor of the soluble PBP antenna that acts as the adaptor to bridge the energy gap and transfer the excitations to the integral membrane photosystems embedded in the thylakoid membrane which have a local absorption maximum at 668 ± 3 nm (Fig. 4de green spectrum).

Having an adaptor protein (*Ha*PE645) coupling the soluble and membrane bound antennas presents an organizational challenge. Once the excitation is transferred to *Ha*PE645 it must then be transferred to a photosystem (or LHC) on the order of a nanosecond (*35*) otherwise it risks energy loss. As the distance of the *Ha*PE645 to the membrane bound antennas increases, the transition probability for excitation transfer between *Ha*PE645 and the membrane system decreases, increasing the likelihood of trapping and fluorescence on *Ha*PE645 (Fig. 4). This implies that for maximal efficiency, *Ha*PE645 should be near the thylakoid membrane which contains both photosystems and integral membrane LHC antennas (Fig. S8; Supplementary Texts G and H). Under the expected range of physiological conditions 50% FRET efficiency is achieved when *Ha*PE645 is either adjacent or better, tethered to the thylakoid membrane.

This implies under normal physiological conditions that *Ha*PE645 would be the closest protein to the membrane (Fig. S8) so as to ensure maximal energy transfer from the bulk antenna to the membrane systems. It may be significant that *Ha*PE645 maintains a distinct *closed* quaternary structure, either as a source of molecular recognition or due to its functional properties. We note that the recent structure of a cryptophyte photosystem I supercomplex has an unknown protein protruding into the lumenal space, potentially providing a membrane-proximal binding site for an adaptor such as *Ha*PE645, potentially orienting it for favorable transfer from the β82 PCB chromophore (*36*). The bulk antenna proteins, *Ha*PE555 and *Ha*PE560, are densely packed to form a concentrated glass of proteins as evidenced by electron micrographs and FRAP experiments. It is also possible that the rafts of *Ha*PE555 filaments seen in all crystal forms are part of the organizing principle of the antenna architecture (Supplementary Text C and H).

Based on this, a picture for light harvesting emerges (Fig. 4f). Sunlight filters through water and visible photons are captured by the bulk antenna proteins *Ha*PE555 and *Ha*PE560 (Fig. 4f, Step 1). Given their spectral properties, energy is transferred readily among these proteins (Fig. 4f, Step 2). Eventually, the excitation is transferred to the *Ha*PE645 adaptor protein (Fig. 4f, Step 3). Within this protein the excitation is transferred to the β82 PCB chromophore on the αSβ protomer (Fig. 4f, Step 3, with panel C showing internal transfer to PCB). From there, the excitation is transferred to the integral membrane photosystems (Fig. 4f, Step 4).

Evidence of a cryptophyte antenna with multiple expressible (and expressed) cryptophyte α chains exists in the literature (*17, 23*) but has been seemingly buried until recently (*24*) (Supplementary Text I). In fact, in organisms with a majority ∼550 nm absorbing PBP, evidence of a 645 nm absorbing protein is present but often discarded as its significance was not fully grasped (*11, 25, 37, 38*). Furthermore, evidence of simultaneous expression of *open* and *closed* forms has been recently discovered in *H. virescens* (*10*).

In *H. andersenii*, we have discovered an energetically complex, multi-subunit antenna comprised of both *closed* and *open* form PBPs, presenting a model for the complete antenna. Given transcriptomics for other cryptophytes (*10*) as well as spectroscopy describing proteins with a 645 nm peak, our model is likely more general for cryptophyte light harvesting, with many cryptophytes having a similar rainbow of proteins. We envisage that a typical cryptophyte soluble antenna would comprise a bulk light harvesting protein with a peppering of other PBPs with specific spectral, structural, and spatial properties so as to energetically couple the soluble antenna to the integral membrane photosystems.

## Acknowledgments

The authors acknowledge use of facilities in the Structural Biology Facility, the Bioanalytical Mass Spectrometry Facility and the Electron Microscope Unit all within the Mark Wainwright Analytical Centre – UNSW. This research was undertaken in part using the MX1 (*39*) and MX2 (*40*) beamlines at the Australian Synchrotron, part of ANSTO, and made use of the Australian Cancer Research Foundation (ACRF) detector. KAM was supported by AOARD (FA2386-17-1-4101). HWR is supported by a scholarship from the Australian Government Research Training Program.

## Funding

Australian Research Council Discovery Project grant DP180103964 (PMGC)

United States Air Force Office of Scientific Research AOARD grant FA2386-17-1-4101 (PMGC)

Zelman Cowen Academic Initiatives (PMGC)

Australian Research Council Linkage Infrastructure, Equipment and Facilities grant LE190100165 (PMGC)

DARPA (QuBE) (BRG)

Natural Sciences and Engineering Research Council of Canada ID 4688 (BRG)

## Author contributions

Conceptualization: HWR, AJL, PMGC

Methodology: HWR, KAM, HI, BRG, PMGC

Investigation: HWR, AJL, KAM, HI, JB, SG, BRG, PMGC

Funding acquisition: BRG, PMGC

Resources: KAM, JB, SG, PT, BRG, PMGC

Supervision: KAM, PT, PMGC

Writing – original draft: HWR, BRG, PMGC

Writing – review & editing: HWR, KAM, HI, BRG, PMGC

## Data and materials availability

All coordinates and structure factors have been deposited with the Protein Data Bank under the accession codes: 8EL3, 8EL4 and 8EL5 -the filament containing structures of *Ha*PE555 from chromatography peak 555A; 8EL6 -the structure of *Ha*PE555 from chromatography peak 555B; 7SSF -*Ha*PE560; and 7SUT -*Ha*PE645.

All other data are available in the main text or the supplementary materials.

## Supplementary Materials

Materials and Methods

Supplementary Text

Figs. S1 to S11

Tables S1 to S5

References(*41*–*59*)

## References and Notes

1. T. L. Richardson, The colorful world of cryptophyte phycobiliproteins. Journal of Plankton Research 44, 806–818 (2022).

2. J. M. Archibald, Cryptomonads. Curr Biol 30, R1114–R1116 (2020).

3. H. W. Rathbone, K. A. Michie, M. J. Landsberg, B. R. Green, P. M. G. Curmi, Scaffolding proteins guide the evolution of algal light harvesting antennas. Nat Commun 12, 1890 (2021).

4. X. You et al., In situ structure of the red algal phycobilisome-PSII-PSI-LHC megacomplex. Nature 616, 199–206 (2023).

5. N. Adir, S. Bar-Zvi, D. Harris, The amazing phycobilisome. Biochim Biophys Acta Bioenerg 1861, 148047 (2020).

6. K. E. Apt, J. L. Collier, A. R. Grossman, Evolution of the phycobiliproteins. J Mol Biol 248, 79–96 (1995).

7. A. B. Doust et al., Developing a structure-function model for the cryptophyte phycoerythrin 545 using ultrahigh resolution crystallography and ultrafast laser spectroscopy. J Mol Biol 344, 135–153 (2004).

8. K. E. Wilk et al., Evolution of a light-harvesting protein by addition of new subunits and rearrangement of conserved elements: crystal structure of a cryptophyte phycoerythrin at 1.63-A resolution. Proc Natl Acad Sci U S A 96, 8901–8906 (1999).

9. S. J. Harrop et al., Single-residue insertion switches the quaternary structure and exciton states of cryptophyte light-harvesting proteins. Proc Natl Acad Sci U S A 111, E2666–2675 (2014).

10. K. A. Michie et al., Molecular structures reveal the origin of spectral variation in cryptophyte light harvesting antenna proteins. Protein Sci 32, e4586 (2023).

11. C. O’hEocha, M. Raftery, Phycoerythrins and phycocyanins of cryptomonads. Nature 184, 1049–1051 (1959).

12. F. T. Haxo, D. C. Fork, Photosynthetically active accessory pigments of cryptomonads. Nature 184, 1051–1052 (1959).

13. M. B. Allen, E. C. Dougherty, L. J. Mc, Chromoprotein pigments of some cryptomonad flagellates. Nature 184, 1047–1049 (1959).

14. A. N. Glazer, G. J. Wedemayer, Cryptomonad biliproteins - An evolutionary perspective. Photosynthesis Research 46, 93–105 (1995).

15. E. Gantt, in Biochemistry and Physiology of Protozoa (Second Edition), M. Levandowsky, S. H. Hutner, Eds. (Academic Press, 1979), pp. 121–137.

16. D. R. A. Hill, K. S. Rowan, The biliproteins of the Cryptophyceae. Phycologia 28, 455–463 (1989).

17. M. J. Broughton, C. J. Howe, R. G. Hiller, Distinctive organization of genes for light-harvesting proteins in the cryptophyte alga Rhodomonas. Gene 369, 72–79 (2006).

18. J. Jenkins, R. G. Hiller, J. Speirs, J. Godovac-Zimmermann, A genomic clone encoding a cryptophyte phycoerythrin alpha-subunit. Evidence for three alpha-subunits and an N-terminal membrane transit sequence. FEBS Lett 273, 191–194 (1990).

19. E. Morschel, W. Wehrmeyer, Multiple forms of phycoerythrin-545 from Cryptomonas maculata. Arch Microbiol 113, 83–89 (1977).

20. E. Morschel, W. Wehrmeyer, Cryptomonad biliprotein: phycocyanin-645 from a Chroomonas species. Arch Microbiol 105, 153–158 (1975).

21. A. N. Glazer, G. Cohen-Bazire, A comparison of cryptophytan phycocyanins. Arch Microbiol 104, 29–32 (1975).

22. C. Brooks, E. Gantt, Comparison of phycoerythrins (542, 566nm) from cryptophycean algae. Arch Mikrobiol 88, 193–204 (1973).

23. B. A. Curtis et al., Algal genomes reveal evolutionary mosaicism and the fate of nucleomorphs. Nature 492, 59–65 (2012).

24. T. Kieselbach, O. Cheregi, B. R. Green, C. Funk, Proteomic analysis of the phycobiliprotein antenna of the cryptophyte alga Guillardia theta cultured under different light intensities. Photosynth Res 135, 149–163 (2018).

25. L. C. Spangler, M. Yu, P. D. Jeffrey, G. D. Scholes, Controllable Phycobilin Modification: An Alternative Photoacclimation Response in Cryptophyte Algae. ACS Cent Sci 8, 340–350 (2022).

26. H. Scheer, K. H. Zhao, Biliprotein maturation: the chromophore attachment. Mol Microbiol 68, 263–276 (2008).

27. K. E. Overkamp et al., Insights into the biosynthesis and assembly of cryptophycean phycobiliproteins. J Biol Chem 289, 26691–26707 (2014).

28. L. K. Anderson, C. M. Toole, A model for early events in the assembly pathway of cyanobacterial phycobilisomes. Mol Microbiol 30, 467–474 (1998).

29. C. Lichtle, Effects of Nitrogen Deficiency and Light of High-Intensity on Cryptomonas-Rufescens (Cryptophyceae) .1. Cell and Photosynthetic Apparatus Transformations and Encystment. Protoplasma 101, 283–299 (1979).

30. M. A. Faust, E. Gantt, Effect of Light-Intensity and Glycerol on Growth, Pigment Composition, and Ultrastructure of Chroomonas Sp. Journal of Phycology 9, 489–495 (1973).

31. E. Gantt, M. R. Edwards, Provasol. L, Chloroplast Structure of Cryptophyceae - Evidence for Phycobiliproteins within Intrathylakoidal Spaces. Journal of Cell Biology 48, 280–& (1971).

32. T. Mirkovic, K. E. Wilk, P. M. Curmi, G. D. Scholes, Phycobiliprotein diffusion in chloroplasts of cryptophyte Rhodomonas CS24. Photosynth Res 100, 7–17 (2009).

33. R. Kana, O. Prasil, C. W. Mullineaux, Immobility of phycobilins in the thylakoid lumen of a cryptophyte suggests that protein diffusion in the lumen is very restricted. FEBS Lett 583, 670–674 (2009).

34. K. M. Heidenreich, T. L. Richardson, Photopigment, Absorption, and Growth Responses of Marine Cryptophytes to Varying Spectral Irradiance. J Phycol 56, 507–520 (2020).

35. C. D. van der Weij-De Wit et al., Phycocyanin Sensitizes both Photosystem I and Photosystem II in Cryptophyte Chroomonas CCMP270 Cells. Biophysical Journal 94, 2423–2433 (2008).

36. L. S. Zhao et al., Structural basis and evolution of the photosystem I-light-harvesting supercomplex of cryptophyte algae. Plant Cell 35, 2449–2463 (2023).

37. C. D. Martin, R. G. Hiller, Subunits and chromophores of a type I phycoerythrin from a Chroomonas sp. (Cryptophyceae). Biochimica et Biophysica Acta (BBA) - General Subjects 923, 88–97 (1987).

38. R. G. Hiller, C. D. Martin, Multiple forms of a type I phycoerythrin from a Chroomonas sp. (Cryptophyceae) varying in subunit composition. Biochim Biophys Acta 923, 98–102 (1987).

39. N. P. Cowieson et al., MX1: a bending-magnet crystallography beamline serving both chemical and macromolecular crystallography communities at the Australian Synchrotron. J Synchrotron Radiat 22, 187–190 (2015).

40. D. Aragao et al., MX2: a high-flux undulator microfocus beamline serving both the chemical and macromolecular crystallography communities at the Australian Synchrotron. J Synchrotron Radiat 25, 885–891 (2018).

41. K. Katoh, K. Misawa, K. Kuma, T. Miyata, MAFFT: a novel method for rapid multiple sequence alignment based on fast Fourier transform. Nucleic Acids Res 30, 3059–3066 (2002).

42. J. J. Almagro Armenteros et al., Detecting sequence signals in targeting peptides using deep learning. Life Sci Alliance 2, e201900429 (2019).

43. D. N. Perkins, D. J. Pappin, D. M. Creasy, J. S. Cottrell, Probability-based protein identification by searching sequence databases using mass spectrometry data. Electrophoresis 20, 3551–3567 (1999).

44. C. C. P. N. 4, The CCP4 suite: programs for protein crystallography. Acta Cryst. D 50, 760–763 (1994).

45. P. D. Adams et al., PHENIX: building new software for automated crystallographic structure determination. Acta Crystallogr D Biol Crystallogr 58, 1948–1954 (2002).

46. P. Emsley, K. Cowtan, Coot: model-building tools for molecular graphics. Acta Crystallogr D Biol Crystallogr 60, 2126–2132 (2004).

47. W. L. DeLano, The PyMOL User’s Manual. (DeLano Scientific, San Carlos, CA, USA, 2002).

48. T. D. Goddard et al., UCSF ChimeraX: Meeting modern challenges in visualization and analysis. Protein Sci 27, 14–25 (2018).

49. C. C. Jumper, I. H. M. van Stokkum, T. Mirkovic, G. D. Scholes, Vibronic Wavepackets and Energy Transfer in Cryptophyte Light-Harvesting Complexes. J Phys Chem B 122, 6328–6340 (2018).

50. K. E. Overkamp et al., Chromophore composition of the phycobiliprotein Cr-PC577 from the cryptophyte Hemiselmis pacifica. Photosynth Res 122, 293–304 (2014).

51. A. J. Laos et al., Cooperative Subunit Refolding of a Light-Harvesting Protein through a Self-Chaperone Mechanism. Angew Chem Int Ed Engl 56, 8384–8388 (2017).

52. V. May, O. Kühn, Charge and Energy Transfer Dynamics in Molecular Systems. (Wiley, 2011).

53. P. M. Krasilnikov, D. V. Zlenko, I. N. Stadnichuk, Rates and pathways of energy migration from the phycobilisome to the photosystem II and to the orange carotenoid protein in cyanobacteria. FEBS Lett 594, 1145–1154 (2020).

54. J. Grabowski, E. Gantt, PHOTOPHYSICAL PROPERTIES OF PHYCOBILIPROTEINS FROM PHYCOBILISOMES: FLUORESCENCE LIFETIMES, QUANTUM YIELDS, AND POLARIZATION SPECTRA. Photochemistry and Photobiology 28, 39–45 (1978).

55. H. W. Rathbone, J. A. Davis, P. M. Curmi, in Photosynthesis in Algae, A. W. Larkum, J. A. Raven, A. Grossman, Eds. (Springer Verlag, 2020).

56. R. MacColl, D. S. Berns, O. Gibbons, Characterization cryptomonad phycoerythrin and phycocyanin. Arch Biochem Biophys 177, 265–275 (1976).

57. A. N. Glazer, G. Cohen-Bazire, R. Y. Stanier, Characterization of phycoerythrin from a Cryptomonas sp. Arch Mikrobiol 80, 1–18 (1971).

58. R. MacColl, W. Habig, D. S. Berns, Characterization of phycocyanin from Chromonas species. J Biol Chem 248, 7080–7086 (1973).

59. C. J. Grisdale, D. R. Smith, J. M. Archibald, Relative Mutation Rates in Nucleomorph-Bearing Algae. Genome Biol Evol 11, 1045–1053 (2019).

