## Supplementary Material for "Molecular dissection of the soluble photosynthetic antenna from a cryptophyte alga"

**This PDF file includes:**

Materials and Methods
Supplementary Text
Figs. S1 to S11
Tables S1 to S5

### Materials and Methods

Transcriptomics sequences

Complete transcriptomes of all four *H. andersenii* strains (MMETSP0043 (strain CCMP644), MMETSP1041 (strain CCMP439), MMETSP1042 (strain CCMP1180), MMETSP1043 (strain CCMP441)) were download from https://www.imicrobe.us/#/projects/104. The transcriptomes were packaged into databases using Genome workbench within which a custom BLAST search was done against published α chain sequences from cryptophyte PBP structures on the PDB (Accession codes: 4LMX, 4LMS, 4LM6, 1XG0). The resulting list of 154 sequences produced was then manually pruned using the known cryptophyte α sequence motif AP-x(9-10)-C to 67 sequences. The mature proteins were then generated using the cut-site motif "AXA" (*18*). There does not appear to be any correlation between the cut site (AxA) and any other mature sequence features. To remove redundancy within the transcriptomes from different strains, we chose the most complete read (longest and least ambiguous) for each sequence across the transcriptomes manually. The resulting list of 22 sequences (Table S1) was aligned using MAFFT (*41*) to a structural alignment of all published cryptophyte α subunits (Fig. 1a).

During this process, one sequence stood out and did not align correctly with the other sequences. This outlier was a tandem sequence comprised of two concatenated α sequences. This transcript read however does have an ambiguous region (denoted by 'X' in the sequence) and in-between the two motifs there exists a predicted 'ADA' cut site which would produce a mature α subunit from the C-terminal copy. In some transcriptomes, this tandem sequence appears to have been split into two separate reads and as such the tandem sequence has been treated as non-biological in this case.

Three peptides ($Ha\alpha^{O}_{10}$, $Ha\alpha^{C}_{7}$, $Ha\alpha^{C}_{8}$) had longer than usual mature N-terminal tails. Furthermore, $Ha\alpha^{C}_{8}$ did not have a discernible AxA cut site, but rather a SFS cut site (as demarcated by TargetP (*42*)). The *closed* form peptides $Ha\alpha^{C}_{7}$ and $Ha\alpha^{C}_{8}$, as well as $Ha\alpha^{O}_{11}$ had longer than usual mature C-terminal tails. In the alignment (Fig. 1a), these N- and C-terminal extensions were truncated but noted with an ellipsis (with the number of residues removed being indicated).

For the analysis of the distribution of the L1 loop in *open* form proteins within the *Hemiselmis* genus, the transcriptomes MMETSP1355 (*Hemiselmis tepida*, Strain CCMP443), MMETSP1357 (*Hemiselmis rufescens*, Strain PCC563) and MMETSP1356 – (*Hemiselmis virescens*, Strain PCC157) were downloaded and the same analysis was performed as above.

*Hemiselmis* *andersenii* cultures

Cultures of *H. andersenii* strain CCMP644 were grown continuously in a temperature range between 23°C and 19°C illuminated by Osram daylight fluorescent tubes with an average radiant power of 1.2 $Wm^{-2}$ at the cultures. Cultures were grown in rehydrated sea salt (Red Sea brand; 33.5 g/L in MilliQ water) with 0.5 mL/L of 2000x concentrated silicate free f/2 trace elements from Algaboost and 0.25 mL/L of 1,000,000x f/2 vitamin mix from Algaboost. Cells were passaged every fortnight into fresh media in a roughly 1:200 ratio of cell inoculant to media. Fresh media was not autoclaved prior to use. Cells were grown in 500 mL batches in 2 L conical flasks with microporous cloth taped to the opening and agitated by lightly swirling every week. Cells were harvested at the same time point as inoculation of new cultures. This was achieved by centrifugation at 2000 g for 40 mins at 4 °C (400 mL centrifuge bottles in a F10BCI-6x500y rotor and Avanti J-E centrifuge from Beckman Coulter). Samples were frozen and stored at -80 °C.

Protein purification

Cell pellets were resuspended using a magnetic stirrer at 4 °C in 50 mL of 25 mM HEPES and 100 mM NaCl buffer at pH 7.5 with a ~ 2 mg of lysozyme, ~50 μg DNAse 1 and one pellet cOmplete EDTA-free Protease Inhibitor Cocktail for 30 mins. The resulting slurry was passed through the Constant Systems cell disruptor three times at a setting of 30 kPa. The homogenized lysate was then sedimented at 48,000 g for 1 hr and 4 °C in an Avanti J-E centrifuge (Beckman Coutler) in the 50 mL tubes with a rotor JA25.50 (Beckman Coulter). The supernatant was mixed in a 1:1 ratio of 100% saturated ammonium sulphate (at 4°C) and stirred for 1 hr at 4 °C. This mixture was sedimented by centrifugation at 40,000 g and 4 °C for 1 hour (400 mL centrifuge bottles in a F10BCI-6x500y rotor and Avanti J-E centrifuge from Beckman Coulter). The supernatant was concentrated using 30 kD Amicon centrifuge concentrators. The sample buffer was exchanged by diluting the concentrated supernatant in a 1:10 ratio with 25 mM HEPES buffer at pH 7.5, then reconcentrating it and repeating this procedure five times.

The concentrated sample was then fractionated using anion exchange chromatography in a HiTrap Q HP 5mL (GE Life Science) column equilibrated in 25 mM HEPES at pH 7.5 eluted using a 0 – 0.35 M NaCl gradient at 1mL/min (Fig. S1, lower chromatogram). Two fractions were separated from this step; a pink fraction with everything that eluted before 110 mL retention volume (including the unbound fraction) and a purple fraction including everything after.

The pink and purple fractions were both independently passed through a cation exchange step in a HiTrap SP HP 5 mL (GE Life Science) equilibrated in 20 mM sodium acetate at pH 5.0 and eluted using a 0 – 0.25M NaCl gradient at 1mL/min (Figs 1b & S1, upper chromatograms). Fractions were extracted as 3-4 mL samples around each absorption peak in each chromatogram.

The largest samples (PE555A, PE555B, PE560A, PE645A) were then purified using size exclusion chromatography using a Superdex GL 200 Increase 10/300 (GE Life Science) column run at 4 °C in 20 mM HEPES, 100 mM NaCl and 1 mM $NaN_3$ at pH 7.5 run over 1.5 column volumes after the column was equilibrated using 1.5 column volumes of the running solution. Resulting samples were finally concentrated using 30 kD Amicon centrifuge concentrators. Samples were frozen and stored in a −80 °C freezer. Samples used for experiments were kept in this buffer unless otherwise stated.

Thylakoid membrane protein purification

Cell pellets were resuspended using a magnetic stirrer at 4 °C in 50 mL of 25 mM HEPES and 100 mM NaCl buffer at pH 7.5 with a ~ 2 mg of lysozyme, ~50 μg DNase I and one pellet cOmplete EDTA-free Protease Inhibitor Cocktail for 35 mins. The resulting slurry was passed through the Constant Systems cell disruptor three times at a setting of 30 kPa. This was centrifugated at 5,000 x g for 10 mins at 4 °C (400 mL centrifuge bottles in a F10BCI-6x500y rotor and Avanti J-E centrifuge from Beckman Coulter). The supernatant was then centrifugated at 12,000 g for 10 mins at 4 °C using the JA.25.50 fixed-angel rotor in the same centrifuge. The pellet containing thylakoid membranes was resuspended in 0.33 M sorbitol, 25 mM Hepes, 20 mM KCl, 5 mM EDTA, at pH 7.8. This was then centrifuged for 30 min at 38,000 g using the same rotor as the previous step. The residual phycobiliproteins were removed by centrifugation of the resuspended pellet two times in the aforementioned buffer and discarding the supernatant. Finally the pellet was dissolved in 0.1% v/v Triton X-100, 0.33 M sorbitol, 25 mM Hepes, 20 mM KCl, 5 mM EDTA, at pH 7.8 so as to solubilize the thylakoid integral membrane proteins. The spectrum was collected using SPECTROstar Nano instrument in a 1 cm quartz cuvette.

Linear absorption spectroscopy

Absorption spectra were taken in quadruplicate for each protein analyzed using the Nanodrop by Thermo Fisher. Spectra were then averaged and normalized to the area under the curve using Mathematica.

LC-MS/MS

An appropriate amount of sample was reduced with 5mM dithiothreitol, 10 mM iodoacetamide followed by tryptic digestion overnight at 37 °C. Digested peptides were desalted with C18 stage tip (Thermo Fisher) before being separated by nano-LC using an Ultimate 3000 HPLC and autosampler system (Dionex, Amsterdam, Netherlands). Samples (2.5 μl) were concentrated and desalted onto a micro C18 precolumn (300 μm x 5 mm, Dionex) with $H_2O$:$CH_3CN$ (98:2, 0.05 % TFA) at 15 μl/min. After a 4 min wash the pre column was switched (Valco 10 port valve, Dionex) into line with a fritless nano column (75μ x ~10cm) containing C18 media (1.9 μ, 120 Å, Dr Maisch, Ammerbuch-Entringen Germany). Peptides were eluted using a linear gradient of $H_2O$:$CH_3CN$ (98:2, 0.1 % formic acid) to $H_2O$:$CH_3CN$ (64:36, 0.1 % formic acid) at 200 nL/min over 30 min. High voltage (2000 V) was applied to low volume tee (Upchurch Scientific) and the column tip positioned ~ 0.5 cm from the heated capillary (T=275°C) of an Orbitrap Velos (Thermo Electron, Bremen, Germany) mass spectrometer. Positive ions were generated by electrospray and the Orbitrap operated in data dependent acquisition mode. A survey scan m/z 350-1750 was acquired in the Orbitrap (Resolution = 30,000 at m/z 400, with an accumulation target value of 1,000,000 ions) with lockmass enabled. Up to the 10 most abundant ions (>4,000 counts) with charge states > +2 were sequentially isolated and fragmented within the linear ion trap using collisionally induced dissociation with an activation q = 0.25 and activation time of 30 ms at a target value of 30,000 ions. M/z ratios selected for MS/ MS were dynamically excluded for 30 seconds. All MS/MS spectra were searched against UniProt and customized database with MASCOT (version 2.3) (*43*) with the following search criteria: enzyme specificity was trypsin; precursor and product ion tolerances were at 4 ppm and ± 0.4 Da, respectively; variable modification of methionine oxidation; and one missed cleavage was allowed. Mass spectrometric analysis was carried out at the Bioanalytical Mass Spectrometry Facility, University of New South Wales, Australia.

Electrospray ionization spectrometry

Samples were dialyzed into water to remove any traces of salt in the sample. Samples were fractionated through reverse phase chromatography on a ACQUITY UPLC Protein BEH C4, 300Å, 1.7 μm, 2.1 mm X 50 mm column (Waters) with a starting buffer of 0.1% formic acid in water (v/v) and final buffer of 0.1% formic acid in acetonitrile (v/v). Electrospray ionization was performed in-line by a Vion QTof mass spectrometer (Waters) using ESI positive sensitivity mode in a mass range of m/z 500-4000 Da. Data were deconvolved using UNIFI (Waters). All equipment is part of the Bioanalytical Mass Spectrometry Facility, Mark Wainwright Analytical Centre, UNSW.

Theoretical masses for all mature α sequences derived from the transcriptome were calculated using *SwissProt* and the mass of a phycoerythrobilin (PEB) chromophore (586.7 Da) was added by hand. Note that some sequences could not be given a mass estimate as they contained either: misreads ('X' in the sequence); they did not contain a start site; or some unknown mature peptides may have a different chromophore bound that is not PEB. When interpreting masses, ladders of 16 Da are observed above the native mass on peptides with multiple methionines, suggesting methionine oxidization. There are mass ambiguities here also given the similarity in size of many α subunits to the resolution of the mass spectrometry setup.

From both the peptide identification by fragment LC-MS/MS and the intact electrospray mass spectrometry, this is by no means an exhaustive analysis. The isolates analyzed were chosen as they were the only ones with a high enough concentration and purity. From chromatograms during purification, it was clear there were further isolates available, but at a concentration and purity that was too low. The mass spectrometry that has been done therefore represents a survey of the most prominent α subunit pairings and was somewhat hampered by cross-contamination between different isolates taken from cation exchange.

Protein abundance calculations

Spectrotype abundance estimates were calculated by first estimating the fraction of pink vs purple proteins and then estimating the subdivisions within each spectrotype. The fraction of pink vs purple was calculated via integration under the curve of the 560 nm absorbance trace in Mathematica and cut at the same site as per purification instructions above. Due to a non-linearity in the 560 nm absorbance arising from saturation of the detector, the chromatogram trace measured at 280 nm absorbance (which was not saturated) was used as a substitute by manually scaling the 280 nm peak (unbound fraction) to the 560 nm absorbance around this peak where the 560 nm signal was small enough to be in the linear regime of the detector. This matching was done by eye. An uncertainty in the scaling was estimated from this procedure based on the limits of a satisfactory fit. Integration of the 560 nm absorbance of the purple fraction was scaled by 7/8 to accommodate the chromophore alteration. The areas under the curve of the pink and purple fractions were compared as a percentage of the total area. Within the pink or purple fractions, the most prominent peaks of the 560 nm absorption in the cation exchange chromatogram for each species (HaPE555A and HaPE560A for pink; HaPE645A for purple) were fit to a Lorentzian-like function (below; with ε a free parameter) using NonlinearModelFit in Mathematica. The area under the curve was then compared to the total area under the curve of the 560 nm absorption to produce a percentage fraction (Table S2). The parameters in the equation below are: $A$ — height scaling, Γ— peak width, $a$ — elution peak position, $x$ — elution volume and $\varepsilon$ — non-Lorentzian factor.

$$\frac{A}{\pi}\frac{\frac{1}{2}\Gamma}{|x-a|^{2+\varepsilon}+\left(\frac{1}{2}\Gamma\right)^2}$$

Excitation-emission maps

Spectra were collected using a Jasco FP-8500 Fluorescence Spectrometer and plots were produced in Mathematica. Truncation of the data at a particular height (for heat maps) was also performed in Mathematica.

Measuring concentrations of PBPs for crystallography

Due to the chromophores, it is not possible to sensibly use UV absorption at 280 nm to estimate protein concentrations for PBPs. As the most stable feature is the optical absorbance from the chromophores, we instead estimate concentration by measuring the absorbance at 560 nm. This provides a useful guide for crystallization experiments, however, the unit is not calibrated in terms of molar concentration.

Crystal structures of *Ha*PE555

*Ha*PE555 fractions were crystallized using a sitting drop and vapor diffusion method in a 96 well plates at room temperature. The crystal for 8EL6 was grown in PEG3350 23.6% (w/v). The crystals for 8EL3-8EL5 were grown in PEG3350 25% (w/v) + AddScreen by Hampton Research. Specifically, PEG3350 25% (w/v) + NaBr 0.01M (8EL3), PEG3350 25% (w/v) + Sarcosine 0.01M (8EL4) and PEG3350 25% (w/v) + Benzamidine HCl 2% (w/v) (8EL5). All drops were 150nL protein and 150nL mother liquor with protein concentration of 0.7 absorbance units at 560 nm with 0.1 mm path length and set by Formulatrix NT-8 and Art Robins Phoenix crystallization robots.

X-ray diffraction datasets were collected at Australian Synchrotron beamlines MX1 and MX2. The experiment number (as part of the Sydney Collaborative Access Program) for 8EL3-8EL5 is 16068g and the experiment number for 8EL6 is 14590a. Data were collected at a wavelength of 0.9537Å (13.0 keV) and temperature 100.0 K. Crystals were cryoprotected in 20% glycerol (8EL3-8EL5) or Paratone-N (8EL6).

For 8EL3-8EL6, derived from peak 555A, data processing for all datasets was performed in DIALS (part of the CCP4 suite; version 7.0.066 (*44*)). Processed data was phased by molecular replacement in phenix.phaser (part of the Phenix suite; version 1.15.2 (*45*)) using the published *H. andersenii* PE555 (PDB 4LMX (*9*)) with all asymmetry removed. All data was reindexed such that the unit cell vector c was pointing along the filament direction. All models were iteratively built using phenix.refine (part of the Phenix suite; versions 1.15.2-1.19.2 (*45*)) and Coot (versions 0.8.9.2-0.9.5 (*46*)).

In the later part of refinement, a non-standard approach had to be taken to resolve the problem of mixed α chains (microheterogeneity). A copy of each α chain (*Ha*$\alpha^{O}{}_{1}$ and *Ha*$\alpha^{O}{}_{2}$) was modelled into each α subunit position as complete alternate conformer chains with different chain names. For residues that had the same identity, the duplicate was removed. The backbone atoms of each pair of residues in each α subunit were restrained to each other in Phenix using a harmonic restraint with a 0 mean and 0.05 σ. During this procedure, all occupancy refinement was turned off and the occupancy of each alternate chain was estimated heuristically using the B-factor of variable residues. Occupancies were manually varied and the most plausible occupancy split was chosen as 50:50 occupancy as this brought the B-factors of each altimer as close as possible. Following phasing and initial model building, we believed that 8EL6 was simply a symmetric dimer of two *Ha*$\alpha^{O}{}_{2}$. Modelling microheterogeneity in the same fashion as 8EL3-8EL5 dispelled the idea that 8EL6 could be symmetric as a 50:50 split seemed to model the density well, not producing any difference density and having equal B-factors. Furthermore, the differences in the C-termini of each alpha chain meant that some waters in the structure had to have occupancies correlated to the alternate chains.

The two α subunits are nearly identical, and the few non-identical residues between the different α subunits do not make any significant crystallographic lattice contacts. As such, the crystal packing in principle cannot differentiate between different arrangements with α chains swapped (i.e. $(\alpha_1\beta).(\alpha_2\beta)$ versus $(\alpha_2\beta).(\alpha_1\beta)$) leading to a statistical mixture of two overlayed chains which could be seen in the electron density. The same problem was encountered in the original PE555 structure (PDB 4LMX (*9*); Fig. S2a and S3a). How can an $(\alpha_1\beta)(\alpha_2\beta)$ pairing take place when there are no non-identical residue pairs on the binding interface? The differences in the C-terminal tails of each α subunit, which do make contact, may provide a mechanism for heterodimerization. It is unclear from the crystallography as to whether *Ha*PE555 is a statistical mixture of $(\alpha_1\beta).(\alpha_2\beta)$, $(\alpha_2\beta).(\alpha_1\beta)$, $(\alpha_1\beta).(\alpha_1\beta)$ and $(\alpha_2\beta).(\alpha_2\beta)$ given the near identity of the α subunits (55/67 residues). However, as noted previously (*9*), it is more likely that each PBP in the crystal is either $(\alpha_1\beta).(\alpha_2\beta)$ or $(\alpha_2\beta).(\alpha_1\beta)$, as the longer $\alpha_1$ would result in a steric clash within a putative $(\alpha_1\beta).(\alpha_1\beta)$ PBP. We note that a similar microheterogeneity has been observed in the recent crystal structure of *Ps*PE545 from *Proteomonas sulcata* where the crystal contains an overlay of $(\alpha_L\beta).(\alpha_S\beta)$ and $(\alpha_S\beta).(\alpha_L\beta)$ (*25*).

In modelling electron density maps, a lot of negative Fo-Fc regions were spotted. This corresponded to larger than van der Waals voids that were present in these structures, as well as *Ha*PE560 and *Ha*PE645. It was thought that these were the result of solvent scaling problems (i.e. Phenix has assumed there is bulk solvent given the size of the voids however, no electron density is observed for that region). Ultimately, this was a minor issue in refinement and ignored. There was also a lot of disorganized solvent density that was put down to partial PEG occupation.

Each α chain hosts a 5-hydroxylysine which can be identified clearly in the density and has been noted previous structures of cryptophyte PBPs (*9*). The identity of each chromophore was also called as either phycoerythrobilin (PEB) or dihydrobiliverdin; (DBV) based on either sp2 or sp3 bonding within pyrrole ring A, which provides clear difference density if a planar ring is swapped for a non-conjugated ring (or vice versa).

Data reduction and refinements statistics for each structure are shown in Table S3.

Crystal structures of *Ha*PE560

*Ha*PE560 was crystallized using a sitting drop and vapor diffusion method in 96 well plates in PEGMME5000 21.4% (w/v) + Tris (pH 6.5) 0.162 M (150 nL protein + 150 nL mother liquor with a protein concentration of 0.7 absorbance units at 560 nm with 0.1 mm path length) at 25°C where drop-setting was performed by the Art Robins Phoenix robot. Crystals were cryoprotected in 10% glycerol. Data collection for the final structure was performed on the Australia Synchrotron Beamline MX1 (experiment number 14590a; part of the Sydney Collaborative Access Program). Data were collected at a wavelength of 0.9537 Å (13.0 keV) and temperature 100.0 K. Image processing was performed in iMosFLM (part of the CCP4 suite; version 7.0.066 (*44*)) to a resolution of 1.45 Å. Processed data was phased by molecular replacement in phenix.phaser (part of the Phenix suite; version 1.15.2 (*45*) using the already published *H. andersenii* β subunit (chain B from PDB 4LMX, i.e. PE555 (*9*)). The final model was iteratively built using phenix.refine (part of the Phenix suite; versions 1.15.2- 1.19.2 (*45*)) and Coot (versions 0.8.9.2-0.9.5 (*46*)).

Data reduction and refinements statistics are provided in Table S3.

Cryptophyte phycobiliproteins usually support post-translational modifications (other than bound chromophores). As with *Ha*PE555, each α chain hosts a 5-hydroxylysine which can be identified clearly in the density and has been noted previous structures of cryptophyte PBPs (*9*). A lot of negative Fo-Fc map regions were spotted throughout modelling and were attributed to the same causes as *Ha*PE555.

One curiosity is that, compared to *Ha*PE555, the entire structure of *Ha*PE560 is dilated along one axis and squeezed along a perpendicular axis, both across the solvent filled hole (*Poisson effect*; Fig. S4). The RMSD between the two β subunits forming the dimer of protomers of *Ha*PE560 and *Ha*PE555 is 0.98 Å across all atoms and 0.25 Å when only a single β subunit from each protein is used. This *Poisson effect* (dilation correlated to an orthogonal contraction) appears to be due to the *L1* loop which nudges the β subunits apart while drawing the α subunits together in a perpendicular direction as it consolidates the open form.

Evidence for the seven-residue loop insertion in the α subunit was clear in the density (Fig. S5a). There were some general regions of the final structure that require care. Firstly, the CD loop of the β subunit is not well ordered but traceable. It is likely that the disorder arises from the fact that the CD loop interacts with a crystallographically neighboring CD loop across a crystallographic two-fold axis. The models for all β chains are also missing the first 2-4 residues as the density was too low to interpret.

At this resolution, some residues began to appear with clearly correlated alternate conformers. Commonly, these were waters, where the alternate conformer of a residue has a water sitting in the primary conformer with occupancy correlated to it. More prominently however, Arg129 of chain F has a secondary conformation which overlaps with a Tris molecule and as such has a correlated occupancy to it. These were all modelled. Some map regions were clearly spaces for more Tris molecules and PEG molecules (in difference density), however, when they were modelled and refined no strong 2Fo-Fc density was present and as such were removed. Ordered solvent was modelled using only the best waters (deleted all waters with e/Å$^3$ <1.5 or B-factor >50 Å$^2$). There were some densities that were evidently a duet of waters with enough conformational freedom to smear them out into an elongated region. Furthermore, there were densities suggestive of polyethylene glycol as have been extensively seen in other structures. These densities however were not modelled as they were rather weak compared to neighboring modelled regions. This structure also has one modelled chloride ion and perhaps contains a few more that were not modelled. The choice to model this density as a chloride was made as the density was rather electron rich for a water molecule (2.876 e/$Å^3$), the distances to neighboring atoms were too great for water hydrogen bonding and the ligands made sense for chloride (bonding to nitrogen atoms).

Lastly, there were some regions where the density was poor. One region is in the β subunit chain B residues 143-146, which is modelled into anisotropic density. The difference density in this region suggests a translated second copy of this chain is present in this region. This displacement was not prominent enough to model and so was left. A further poor region is residues 28-33 of the α subunit which always seems poor. This region has few constraints and sits atop the α chain chromophore and is solvent exposed. Chain A (α subunit) Asn29 appears to have an alternate conformer, however, due to the disorder of this loop was not modelled. Residue Phe30 of each β subunit, also has poor map-model correlation. This residue makes no crystallographic contacts and is on the surface of the protein and thus disordered.

Crystal structures of *Ha*PE645

*Ha*PE645A was crystallized using a sitting drop and vapor diffusion method in 96-well plates and in condition PEG3350 25% (w/v) + 0.1M Bis-Tris (pH 5.5) (150nL protein + 150nL mother liquor drops with a protein concentration of 0.32 absorbance units at 560nm with a path length of 0.1 mm) at room temperature. With drop-setting by Formulatrix NT-8 crystallization robot. All gradient optimization trays were produced using the Starlet liquid handling robot from Hamilton. Crystals were cryoprotected with Paratone-N.

Data collection for the final structure was performed on the Australia Synchrotron Beamline MX1 (experiment number 16068e; part of the Sydney Collaborative Access Program). Data were collected at a wavelength of 0.9537 Å (13.0 keV) and temperature 100.0 K. Data processing was performed in DIALS (part of the CCP4 suite; version 7.0.066 (*44*)) to a resolution of 1.49 Å. Processed data was phased by molecular replacement in phenix.phaser (part of the Phenix suite; version 1.15.2 (*45*) using the already published *H. andersenii* β subunit (chain B from PDB 4LMX, i.e. PE555 (*9*)). The final model was iteratively built using phenix.refine (part of the Phenix suite; versions 1.15.2- 1.19.2 (*45*)) and Coot (versions 0.8.9.2-0.9.5 (*46*)).

Data reduction and refinements statistics are provided in Table S3.

A lot of negative Fo-Fc map regions were spotted through-out modelling and were attributed to the same causes as *Ha*PE555 and *Ha*PE560.

As with *Ha*PE560, there are a few alternate conformers with waters in them. These were not modelled (α subunit Chain A Arg45 and Lys64, β subunit Chain B DBV201 and α subunit Chain E Lys64). There were many high B-factor, anisotropic waters evident in the density. These were largely disregarded as they were low confidence and may have been confounded by low occupancy PEG. The same procedure for water picking was used as with *Ha*PE560. As with *Ha*PE560, a single chloride was also modelled and given the same justification. There were many densities indicative of PEG. Most of these were not modelled as they were either broken, branched or weak. These were however clear in the maps and an attempt was made to model them with only a few being left modelled. PEG molecule (Residue 1 of chain I — PG4 surrounding α subunit chain A Lys78) has some problems in its geometry but is physically reasonable and possibly has a water above the NZ nitrogen (not shown). Another PG4 molecule surrounding a water molecule that is hydrogen bonded to the side chain of Asn138 of the β subunit is shown in Fig. S5c. There were at least 2 other densities suggestive of Bis-TRIS that were not modelled as such. One high-confidence molecule was modelled and kept. Others were modelled but later removed as there was too much conformational freedom present creating more problems. Some alternate conformers (which were in weak density) were not modelled as they generated clashes. This is largely since the alternate conformers were correlated. Many of these were not modelled as they were minor. Examples include β subunit chain F Ser55 with Met134, β subunit chain F Ile9 with Thr95. As with *Ha*PE560, the N-terminus of the β subunit was disordered (residues 1-15) and was not modelled. The Phe30 of each β subunit, unlike some other structures is not disordered and packs with neighboring Phe30 residues. Phe30 has been seen to interact with neighboring Phe30 residues in the crystals of both *open* and *closed* forms. Lastly one β82 chromophore of each PBP was a PCB instead of a PEB. The density clearly showed a sp2 bonding between the two rings C and D (Fig. 3b). PEB was initially fit to test if biasing the phases would leave the identity ambiguous, however it was clear in the difference density that it could not be PEB.

Chromophore dihedral angle analysis

Although largely conjugated, the linear tetrapyrrole chromophores deviate from coplanarity between adjacent pyrrole rings due to strain induced by protruding methyl groups. The central pair of rings (B and C) are, in most cases, coplanar at the resolution of the crystal structures and were not analyzed in detail. However, the two outer pyrrole rings (A and D) tend to be twisted with respect to the central pair. To analyze this twisting, we have defined a set of dihedral angles as described in detail in (*10*). Briefly, the coordinates for each chromophore were passed into Mathematica and planes were fit to each of the four pyrrole rings. The molecular geometry can be described by a pair of dihedral angles ($\theta_{inner}$, $\theta_{outer}$) starting from the central pyrrole.

Electron microscopy

1 mL samples of *H. andersenii* cells for electron microscopy were taken at the peak of the growth cycle (14 days post inoculation) and prepared by sedimentation at 1000 g for 5 mins in a benchtop microfuge and resuspended to a volume of 250 µL in MilliQ water. To achieve near-to-native state ultrastructure, cells were pelleted at 1000 x g, loaded onto 6 mm gold coated copper high-pressure freezing planchettes (Leica Microsystems) and high-pressure frozen using a Leica EM ICE (Leica Microsystems). Samples were stored in liquid nitrogen and transferred to the automated freeze substitution apparatus Leica EM AFS (Leica Microsystems), containing a solution of 1% osmium tetroxide, 0.2% uranyl acetate (w/v) and 10% water in acetone. Samples were kept at −90 ºC for 48 h, slowly warmed to −80 ºC (5 ºC /h), kept at that temperature for 3 h, warmed to and kept in the same way at −60 ºC, − 40 ºC and −20 ºC and finally warmed to 0ºC (5 ºC /h). Two washing steps with cold acetone were carried out and the cells were infiltrated overnight with increasing concentrations of Procure resin at room temperature. After 2 changes into fresh Procure resin, the samples were polymerized at 60 ºC for 48 hours. 70 nm sections were cut with a diamond knife and collected onto carbon coated copper slot grids and post stained with 2% uranyl acetate and lead citrate. Grids were imaged using a Jeol 1400 transmission electron microscope (Tokyo, Japan) operating at 100kV. To determine the separation between thylakoid membranes, distances between consecutive thylakoid membranes were approximated by using an edge detection algorithm. Images were imported into Mathematica, cropped to show sections with defined striations and aligned so the striations were completely vertical. Using the inbuilt function EdgeDetect and then calculating the distances between consecutive edges, a histogram of distances was generated. From this histogram a mean and standard deviation was produced.

Fluorescence spectra

Fluorescence spectra were produced by summing each excitation-emission map data set along the excitation axis to get the total fluorescence. The Rayleigh scattering peaks were removed from the integrated fluorescence spectra by fitting and subtracting off a $\propto r^n$ function to the spectrum using NonlinearModelFit in Mathematica.

All spectra were normalized so the maximum intensity was unity. Graphs were also plotted in Mathematica.

Protein analysis and graphics

Protein dimensions were estimated using Pymol (*47*).

Buried surface area was measured in ChimeraX using the function *measure buriedarea*. This included chromophores but no solvent.

All images of protein structures were generated either in PyMol (*47*) or in ChimeraX (*48*).

**Supplementary Text**

A—Unclassified peaks in chromatography

Three peaks in the chromatography did not appear to fit a specific spectrotype when examining their absorption spectra (Fig. S10). These were labelled XA, XB and XC. Each spectrum had features of multiple spectrotypes indicating that they are mixtures possibly due to lack of resolution between peaks or contamination from larger neighboring peaks. Furthermore, the mass spectrometry for fraction XA has, amongst masses for *Ha*PE555 type proteins, a mass for $HA\alpha^{C}{}_{1}$/$HA\alpha^{C}{}_{2}$ and the spectrum has a minor peak at 645 nm suggesting that a *closed* form (*Ha*PE645 type) protein makes up some of the protein components within this peak and producing the small hump at 645 nm. Mass spectrometry shows that *Ha*$\alpha^{OB}{}_{1}$ is also seen in the purple fraction suggesting there is some contamination arising from when pink and purple were split after the initial anion exchange chromatography.

B—*Ha*PE555 from the smaller peak 555B

In all *Ha*PE555 structures, we observed Phe30 of the β subunit making close contacts between adjacent filaments forming 2D sheets (Fig. S3). In the structure derived from chromatography peak 555B (8EL6; Fig. 1b), there is an alteration in the secondary structure of the β subunit around Phe30 with a lengthening helix hA and shortening hY altering the interfilament interaction (Fig. S3bc). This reorganization of secondary structure has not been observed in any other structure of a cryptophyte PBP. This alteration is only observed for one of the two β subunits in this PBP. Given that this peak 555B separates from 555A, this suggests that this alteration in secondary structure is stable and possibly the cause of this separation. This may therefore imply that chromatography peak 555C has this alteration on both β-subunits.

C— *Ha*PE555 filaments

All five crystal forms of protein *Ha*PE555 are constructed from continuous filaments of PBPs (Fig. S2b). The number of PBPs per asymmetric unit varies, either one (8EL4 and 8EL6), two (8EL3 and 8EL5) or three (4LMX). The filament structures observed in the five different crystals are generated from a knob and hole packing but differ from each other in the distances between adjacent molecules along the filament direction (having either tight or loose interfaces; Fig. S2b; Table S4). The knob and hole contact is formed by the CD-loop of one β subunit slotting into a hole formed around the GH loop on an opposite β subunit (Fig. S2b, left and right panels for tight and loose interfaces, respectively). The cleft along this interface tightens in different conditions, where a layer of waters is removed in each step of shortening. The packing changes along the filament direction are evaluated by the average period along the filament, |**c**|/molecule, which is measured by length of the unit cell vector, |**c**| (which is parallel to the filament axis), divided by number of molecules in the ASU (which has been chosen to lie along the filament). The lower the values of |**c**|/molecule, the tighter the packing along the filament axis (Table S4).

From these data alone, it remains unclear if these filaments are biologically relevant, however, two possible effects are apparent. The first is simply that filaments provides a means of organization by clustering together light harvesting units of a single type. The second is that by creating a filament, an organism effectively brings chromophores from neighboring proteins as close as they would be within a single protein thereby increasing excitation energy transfer rates (Fig. S2c). Some chromophores between complexes in a filament are much closer than those intra-protein. Distances between the center of each chromophore in all structures have been calculated (Fig. S2de), along with those of adjacent molecules in *Ha*PE555 filaments (Fig. 2c). Creating a filament also generates spatial order between chromophores of neighboring PBPs. The alignment of dipoles between chromophores maximizes the FRET rate between them and the size of the cluster of chromophores that are coherent with one another.

We note that this filament arrangement in not general for *open* form PBPs as no other *open* form PBP structure to date has been observed to form filaments in this manner and it is unknown if filaments form *in vivo*. Additionally, the crystal structure of *Ha*PE560 does not display filaments. Here the crystal is formed from sheets with two $(\alpha\beta)_2$ molecules in the ASU.

To explore the possibility of filaments in *open* forms other than *Ha*PE555, we generated pseudo-filaments based on *Ha*PE555 structures by performing a least squares structural alignment on multiple copies of the *open* form of interest (Fig. S9). The comparison of all *open* form pseudo-filaments showed that the few residues preceding the β strand S2 in each α chain are critical to the formation of filaments as, in the case of *Ha*PE560, this caused a steric clash with helix hFb of the adjacent molecule in the filament (next to the β-82 chromophore). In this region, *Ha*PE560 forms a single turn helix that protrudes from the surface (Fig. S9, magnified view on right, *Ha*PE560 in purple) whereas *Hv*PC612 (red) and *Hp*PC577 (cyan) have a smaller protrusion which closely follows the adjacent N-terminal end of helix hY (Fig. S9, magnified view, helix on lower right) with a greater capacity for the formation of filaments. *Ha*PE555 (Fig. S9, green) does neither, instead following closely along the β strand S2 (which is likely why we only see tight filaments in the crystals of *Ha*PE555).

D—*L1* loop insertion in *Hemiselmis* α subunits between β strand *S2* and the α helix

Multiple sequence alignment of *Hemiselmis* α subunit sequences from transcriptomes has identified two clusters of sequences that possess an insertion between β strand *S2* and the α helix (Figs 1a, 2f and S7). Ten of these sequences correspond to the *open-braced* structure described in the main manuscript (Fig. S7 labelled canonical *open-braced*, OB +1 and OB +3). In these sequences, the *L1* loop insertion starts after an anchoring aromatic residue (tyrosine or phenylalanine) after β strand *S2* and terminates at an extra N-terminal turn on the α helix (RIG motif). The *L1 open-braced* motif is:

$$G\text{-}x\text{-}x\text{-}G\text{-}x_{1\text{-}3}\text{-}D\text{-}R$$

Ten sequences conform with this motif (clusters labelled canonical *open-braced*, OB +1 and OB +3). We note that prior to the N-terminal glycine, there is a conserved aromatic residue (phenylalanine or tyrosine) which anchors the *L1* loop.

Furthermore, there are additional sequences that contain a similar, but distinct, insertion in the same position (Fig. S7 clusters labelled OB -2 and OB -4). These sequences come from the same organisms which contain the *open-braced* form: *H. andersenii*, *H. rufescens* and *H. tepida* each containing two sequences with these shorter insertions. Most sequences in this cluster (bar $Hr\alpha^{O}$ in cluster OB -4 from *H. rufescens*) contain the motif:

$$G\text{-}x_{1\text{-}2}\text{-}G\text{-}x\text{-}G$$

We note that one of the intact PBPs identified by mass spectrometry contains two of these sequences: $Ha\alpha^{O}_{3}$ and $Ha\alpha^{O}_{4}$ (Fig. 1a and 1e) and it belongs to peak XA in Fig. 1b. The structure of the *L1* insertion in this protein will have to await further investigations.

The alignment of all *open* form and *open-braced* form sequences highlight another conserved feature. At the end of the α helix, there is a conserved Asn-Tyr motif (Fig. S7). This aromatic residue marks the end of the α helix, anchoring the subsequent C-terminal loop. Finally, we note that two sequences in Fig. S7 contain a single residue insertion just prior to the characteristic Asp insertion two residues before the cysteine covalent chromophore attachment site (Fig. S6, arrow just after β strand *S1*). Given that the Asp insertion is responsible for the transition between the *open* and *closed* quaternary structure, it is unclear as to what this extra insertion may create in terms of protein structure.

E—Chromophore geometry and energy transfer relevant parameters

The energetic properties of each chromophore are determined by its identity, geometry, and local electrostatic environment (*10*). *Ha*PE555 and *Ha*PE560 have identical chromophores (DBV on β50/61 and PEB on all other sites) whereas *Ha*PE645 differs on two points (DBV on the α subunits and the single PCB on the β82 site in the $\alpha_2\beta$ protomer; Table S5). The geometry of the three β subunit chromophores (save the β82 of the $\alpha_2\beta$ protomer of *Ha*PE645) is largely identical between these structures within the protein matrix given they all have the same β subunit.

The key differences in chromophore geometry are those for β82 in *Ha*PE645 and the α chromophore in *Ha*PE560, both discussed in the main text, with subtler differences revealed when comparing the chromophore dihedral angles (Fig. S6). For the α chromophore, the dihedral angles between pyrrole rings A and B partition into *closed* and *open* form clusters (Fig. S6a-c) where the major change is due to the insertion of the aspartic acid in the sequence just before the cysteine residue covalently attached to the chromophore. In the *open-braced* form, the *L1* loop shifts and rotates the chromophore (Fig. 2de) which is reflected in the dihedral plot (Fig. S6c). It is likely that the changes in the α chromophore are responsible for the spectral shift between *Ha*PE555 and *Ha*PE560.

For the *closed* form *Ha*PE645, the key difference is the switching of chromophores attached to cysteine β82. The geometry of the PCB chromophore attached to Cys-β82 on the $\alpha_S\beta$ is similar to that observed for *Cs*PC645 from *Chroomonas* sp (Fig. 3de). In terms of chromophore dihedral angles for pyrrole rings C and D, this chromophore forms a cluster with the two β82 chromophores from *Cs*PC645 (Fig. S6de).

The most prominent difference between the protein matrices themselves is between the *closed* form *Ha*PE645 and the two *open* forms. In *closed* forms, the β50/61 chromophores from opposite β subunits are brought into van der Waals contact (Fig. S2e, circled in lower panel), creating a strongly coupled quantum system, which is expected to broaden the spectrum. Strikingly, however, the shape of the main absorption peak is much the same between all three proteins (Fig. 1c; apart from the blue shift in *Ha*PE555 compared to the other two proteins). It is unclear why so little spectral broadening is observed for the *closed* form *Ha*PE645. As for the purpose of *closed* forms then, there is some suggestion that the tight binding of the central chromophores contributes to the enhanced excitation energy transfer rate (by a factor of x2 (*49*)), which may be a benefit of having *closed* form as the terminal acceptor of the soluble antenna and adaptor to the integral membrane systems.

F—Mechanisms for chromophore alterations in *Ha*PE645

The asymmetry in the identity of the β82 chromophore, with PEB attached to $\alpha_L\beta$ versus PCB attached to $\alpha_S\beta$, raises question as to how this asymmetry is achieved on otherwise symmetric β subunits. No such asymmetry has been observed before in cryptophyte PBPs. The current model for how chromophores are covalently attached by lyases in cryptophytes (*50*) is based on an earlier model for phycobilisome assembly (*28*). In these models, lyase enzymes covalently attach linear tetrapyrrole chromophores to specific cysteine residues in the plastid stroma. Chromophore attachment is coupled to folding and assembly of mature proteins prior to transit into the thylakoid lumen. These processes may involve molecular chaperones and folding/degradation pathways that control the quality and integrity of mature light harvesting proteins. Individual lyases are likely to bind to folded and/or partially assembled subunits to recognize the correct cysteine to ensure that the correct chromophore is attached.

For the β82 chromophore, the only distinguishing signal comes from the α subunit associated that forms the αβ protomer. The β82 chromophore site lies proximal to the N-terminus of the α subunit. For *Ha*PE645, the two α subunits have distinct N-terminal sequences in the vicinity of the β82 chromophore. Thus, it is likely that the lyases responsible for loading the two distinct β82 chromophores bind to the assembled αβ protomer (or possibly the fully assembled $\alpha_1\beta.\alpha_2\beta$ complex) so as to attach the correct chromophore. It has been shown that the α subunit itself acts as a chaperone for the assembly and stability of the β subunit (*51*). Finally, we note that the $\alpha_L$ subunit would still have a thylakoid lumenal targeting sequence extending its N-terminus while it resides in the stroma. This targeting sequence may confer a signal to the lyase.

G—FRET efficiency for the model of the *H. andersenii* antenna sandwiched between the thylakoid membranes

A model for the *H. andersenii* antenna is presented in Fig. 4. In this model, the soluble light harvesting proteins are sandwiched between the thylakoid membranes. The proteins are close packed and lack long range mobility. The bulk of the antenna is composed of *Ha*PE555 plus *Ha*PE560, which capture the most energetic photons. The question is: what is the location of the adaptor, *Ha*PE645, that accepts energy from the bulk of the antenna and transfers it to the integral membrane photosystems. To understand the constraints on the location of the adaptor, we use the model to calculate the effect of adaptor location with respect to the membrane on the efficiency of light harvesting.

FRET efficiencies were calculated analytically using a model that assumes all energy captured by the soluble PBP antenna eventually reaches the adaptor, *Ha*PE645, from which it is then transferred via FRET to the integral membrane photosystem, PS-II or others. The model begins with the FRET efficiency equations (*52*) where $k_{ET}$ is the sum of FRET rates to all acceptors and $k_F$ is the rate of fluorescence. $k_{ET}$ itself is given by the equation below where $R_0$ is the Förster distance and $r_i$ is the distance to the i$^{th}$ acceptor. It is assumed that $R_0$ is the same for all donor acceptor pairs (*Ha*PE645—PS-II/Others).

$$E = \frac{k_{ET}}{k_{ET} + k_F}$$

$$k_{ET} = k_F \sum_i \left(\frac{R_0}{r_i}\right)^6$$

$$E = \frac{1}{1 + \left(\sum_i \left(\frac{R_0}{r_i}\right)^6\right)^{-1}}$$

The approximation in this model begins by assuming that *Ha*PE645 has a membrane acceptor directly above and below it. This leads to the equation below.

$$E = \frac{1}{1 + \left(R_0{}^6 \left(\frac{1}{(z-L)^6} + \frac{1}{(z+L)^6}\right)\right)^{-1}}$$

The equation for FRET efficiency as a function of position in the thylakoid ($z$) is given by the equation above where L is the half-width of the thylakoid lumenal space (distance between the bounding membranes).

The FRET efficiency, $E(z)$, was plotted for a range of values of $R_0$ and $L$ (Fig. S8). The width of the lumen (2L) is also taken to be either 12.7 nm (Fig. S8a) or 25 nm (Fig. S8b) and $R_0$ is taken to be in a range of 3 - 6 nm (*53, 54*). From the plot, it can be seen that efficiency drops of quite rapidly and for maximal efficiency, the *Ha*PE645 should be placed near or tethered to the membrane.

H—Model including filaments

The crystal structures of *Ha*PE555 indicate that some cryptophyte PBPs have a propensity to form filaments. It is possible that filament formation occurs in the biological antenna allowing for segregation of different components (to keep the *Ha*PE645 near the membrane). Another possibility is that order and regulation within the thylakoid lumen is generated by tethering the adaptor through another protein or electrostatically as a variety of isoelectric points is observed in the phycobiliproteins of other cryptophyte species (*19*-*22*). Regulation could be triggered by a pH decrease in the thylakoid lumen from increased photosynthesis, releasing tethered *Ha*PE645 from the membrane to slow energy transfer and release oxidative stress.

In a model that includes filaments (Fig. S11), light capture by cryptophytes is likely to proceed as follows. The model begins with the illumination of a cryptophyte by sunlight. A single solar photon with a particular wavelength (or range) sends all chromophores excitable by this wavelength into a quantum superposition. On a sub-femtosecond time scale, this quantum superposition collapses and the photon sends a chromophore or small coherent cluster of chromophores in a superposition to an excited state. Given the large fraction of *Ha*PE555 in the antenna, the excitation most likely exists as a coherent cluster on *Ha*PE555. This coherent cluster explores interactions with neighboring chromophores and the surrounding protein, expanding and/or localizing the cluster, respectively. The excitation may transfer to other chromophores/clusters incoherently via FRET along its sojourn. The coherent cluster present on *Ha*PE555 is likely to be only 2-3 chromophores in size. As the energy landscape of the chromophores of *Ha*PE555, or indeed filaments composed of *Ha*PE555, is roughly flat (because PEB and DBV have similar excitation peaks), the coherent cluster is free to diffuse along this filament. This is also the case for free *Ha*PE560 complexes. At some stage after travelling up to tens of nanometers, the coherent cluster interacts with a nearby *Ha*PE645 onto which it transfers. Once the excitation is present on *Ha*PE645 it migrates around the protein exploring the single quantum structure formed by the central pair of doubly-linked DBV β50/61 chromophores. From this position the excitation can then interact with and migrate to the single β82 PCB. This, now lower energy excitation, can interact with the integral membrane light harvesting complexes comprised mainly of carotenoids and chlorophylls, to which it will transfer (presumably by incoherent FRET) and migrate finally to the special pair of chlorophylls in the photosystem where ultimately, charge separation occurs. This whole process must take place on a nanosecond timescale otherwise the excitation will escape via fluorescence (*55*).

I—History: The path to dissecting the cryptophyte antenna

Evidence for a multi-component antenna is present in early papers describing the soluble light harvesting systems of cryptophytes mainly expressing phycoerythrins. Early absorption spectra of aqueous extracts show that although the soluble antenna protein have predominant maxima around 545nm or 560-568 nm, they have smaller peaks or shoulders around 600-650nm that are somewhat variable. These studies include: *Cryptomonas ovata* (*12, 13, 56*), *Sennia* sp (*Hemiselmis parvula*?) (*11*), *Rhodomonas* CS24 (*37*), and *Cryptomonas acuta* (*16*). Regarding *Rhodomonas* CS24, Martin and Hiller (1987) stated: "It is suggested that one or more of the α subunits of this phycoerythrin may provide the intermediate components necessary for energy transfer to chlorophyll a." (*37*). It is likely that some of this wisdom was lost as protein preparations improved and laboratories focused on the most abundant components.

In addition to the spectroscopic evidence, there was abundant evidence from isoelectric focusing studies that each cryptophyte expressed multiple α subunits with different pIs. These include: *Cryptomonas* sp (*57*), *Cryptomonas maculata* (*19*), *Cryptomonas ovata* (*22*), *Rhodomonas* (*22*), *Rhodomonas* CS24 (*38*), *Chroomonas* (*20, 58*) and *Hemiselmis virescens* (*21*).

J—Evolutionary perspective and α sequences from other *Hemiselmids*

What is curious is why there is such a diversity of α subunit sequences in *H. andersenii*, yet most of the light harvesting appears to be done by one protein and furthermore many of the other proteins that are expressed have similar spectral properties. It is possible that the levels of all of these PBPs are regulated by environmental conditions such as light color, intensity and pH of the lumen. A recent study of photoacclimation in cryptophytes has shown that the spectral properties of isolated PBPs change when the organisms are grown in spectrally altered light (*25*). Given the diversity of α subunit peptides, the levels of these mature light harvesting proteins are likely to be regulated in the cell nucleus by controlling α subunit gene expression.

One thing we do not understand is whether there is a possibility of mixing-and-matching α subunits to produce a statistical mixture of PBPs – however, cells may either have mechanisms to prevent this (such as specific chaperones) or such mixing may simply not occur due to steric selectivity. In the process of PBP maturation, it is likely that αβ protomers fold and assemble independently, and then bind to other αβ protomers rather than folding concurrently as a full set of four chains (*17, 51*). As such, it may be that molecular recognition between αβ protomers selects for specific pairing from the surface generated by the α subunit resulting in the well-defined mature proteins that we have observed in crystal structures.

One further observation is the diversity of α subunits in cryptophyte antennas compared to the shear lack of sequence diversity in β subunits across species (*3*); the progenitor α subunit genes were relocated to the nucleus from the red algal nucleus following secondary endosymbiosis where progeny α subunits proliferated while the β subunit gene remained a single, chloroplastic gene. In fact, nuclear genes have roughly 3x the mutation rate of plastid genes in *H. andersenii* (*59*). The implication for evolution here may be that α subunits are free to diversify and roam the sequence space, finding new possible niches while the β subunits remain relatively constant. The diversity of α subunits is key to an apparent modularity in cryptophyte light harvesting and its potential evolution, adaptation and tunability.

In an evolutionary context, *H. andersenii* highlights the diversification of light harvesting in cryptophytes. Firstly, it presents a mixture of *open* and *closed* forms as well as the novel *open-braced* form. Evidence for the *open-braced* form (including the shorter version of the *L1* loop) has been observed in a handful of *Hemiselmis* transcriptomes (*H. andersenii*, *Hemiselmis tepida* and *Hemiselmis rufecens*) where the *L1* loop insertion is seen (Fig. 2 and S7). This is in part due to limited data, however, it may reflect evolutionary history and a functional role for the *open-braced* form in these organisms. As noted in the main text, although the *open-braced* form is likely more stable than the *open* form, these two forms co-exist in these organisms and in fact in the lumen of the thylakoid, the *open* form remains the dominant protein by mass. Since the innovation of the *open-braced* form is observed in a handful of *Hemiselmis* species, it is possible that *H. andersenii*, *H. rufecens* and *H. tepida* represent the most recent evolutionary branch (given current data) in the cryptophytes, with the development of the *open* forms more generally arising in *Hemiselmis*. Finally, we note that two α subunit sequences from *H. rufecens* have an extra single amino acid insertion before the aspartic acid insertion that differentiates the *open* form, forming an entirely new innovation. It is unclear what this ‘double-open’ insert is doing to the structure.

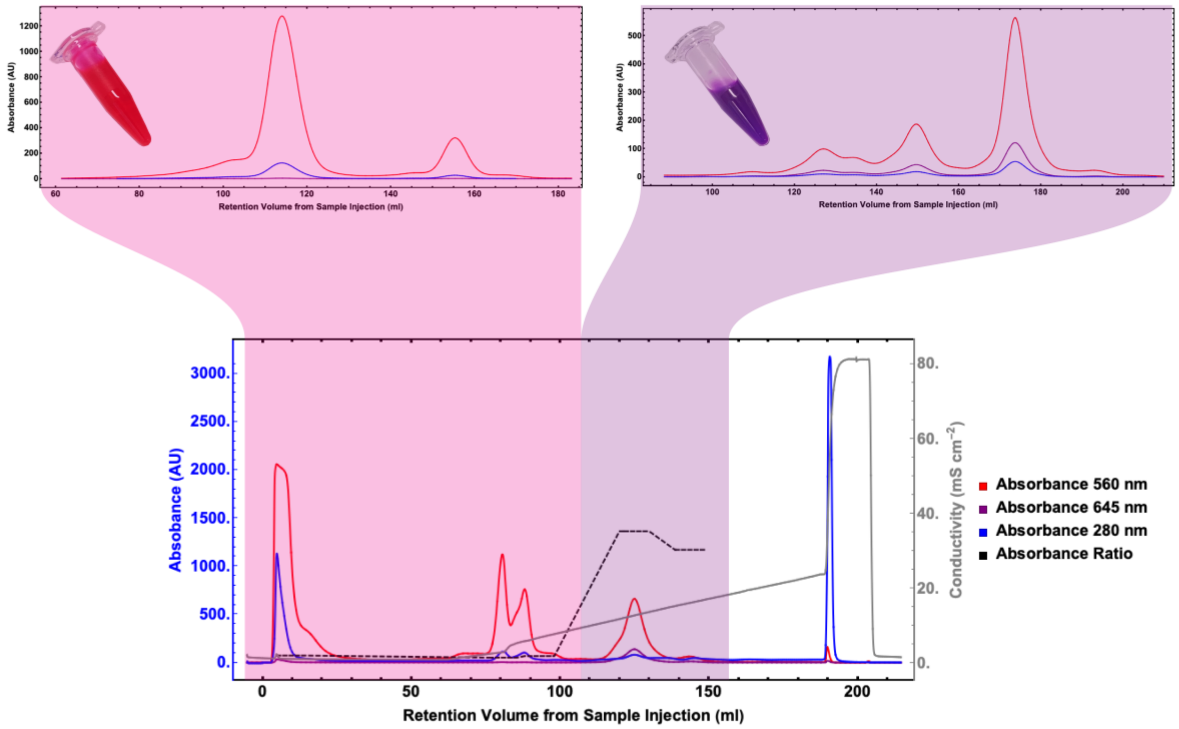


**Figure S1. Purification of soluble light harvesting protein.** Lower chromatogram shows the separation of soluble light harvesting protein into pink and purple fractions using anion exchange chromatography. The two upper panels show further purification of each of these pooled fractions by cation exchange chromatography.

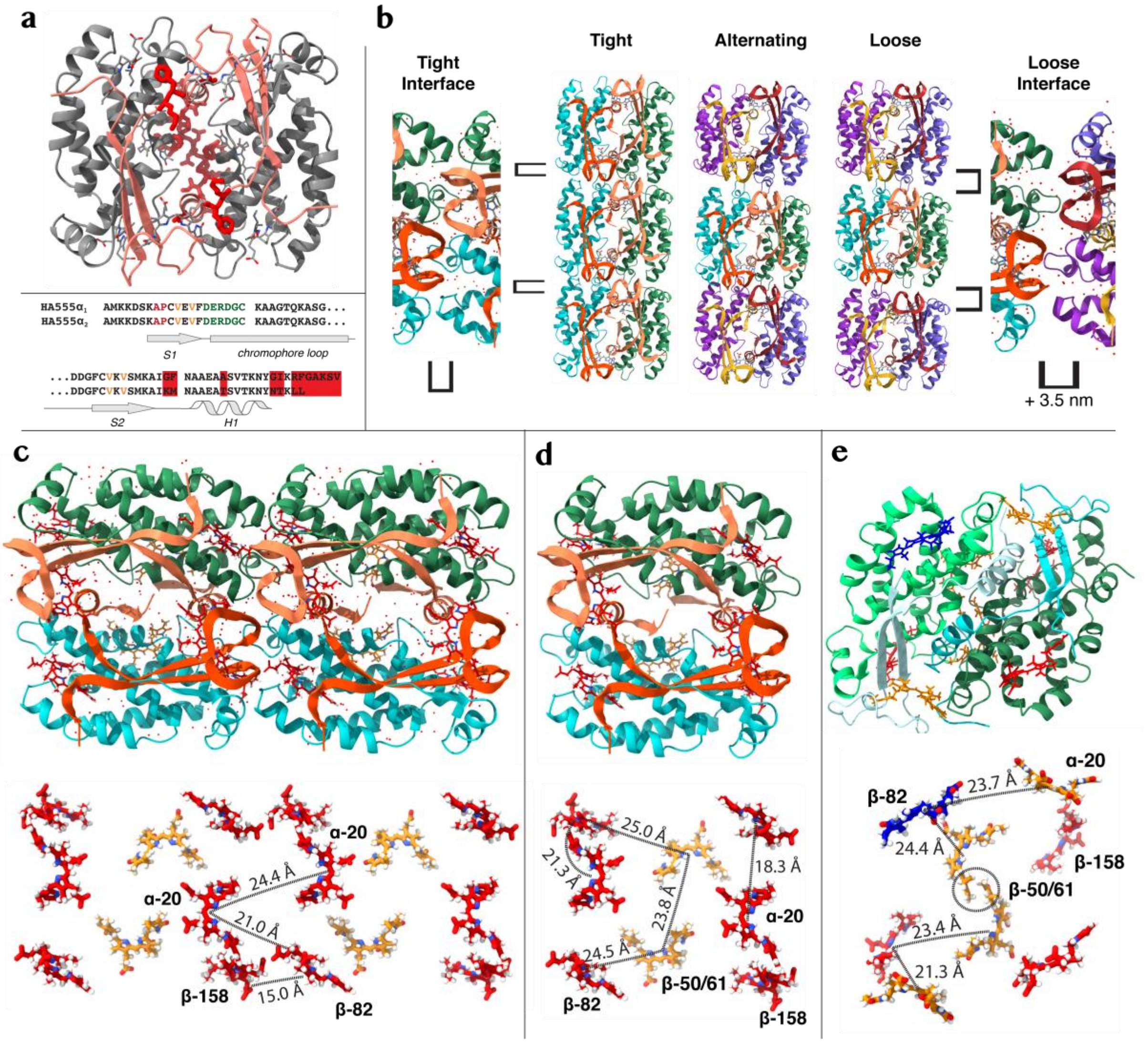


**Figure S2. The crystal structures of *Ha*PE555 are composed of sheets of filaments. a.** The crystal structure of HaPE555 shows a quasi-symmetric dimer with an *open* form quaternary structure which is identical to previously published 4LMX. The two α subunits (peach ribbon) show only minor sequence differences (highlighted in red as overlapped stick figures on the structure, top, and the sequence, bottom). These minor sequence differences result in microheterogeneity in all crystal forms, hence each site is a 50:50 mix of the two distinct residues. **b.** Each crystal form of *Ha*PE555 is composed of 2D layers of *Ha*PE555 filaments that differ in the packing of the interface between monomers along the filament axis. The left most image is the tight interface while the right most image is the loose interface (each shown rotated 90° with respect to the filament images in the central panel). The central part of the panel shows the three distinct filament types, where the left most filament shows tight interface packing, the right most filament shows loose interface packing and the central filament shows alternating tight-loose packing. The nature of the interfaces is indicated by the square brackets, where loose interface has a 3.5 Å wide layer of ordered solvent between two neighboring proteins. The presence of filaments in all crystal forms may have biological significance. **c**-**e**. Show the distances between chromophores for: **c**. the interfilament tight interface; **d**. the intramolecular distances for *Ha*PE555; and **e**. *Ha*PE645. Chromophores are shown as sticks with carbon atoms colored: red – PEB; orange – DBV; and blue – PCB. The upper panels show the complete structures that support these chromophore arrangements.

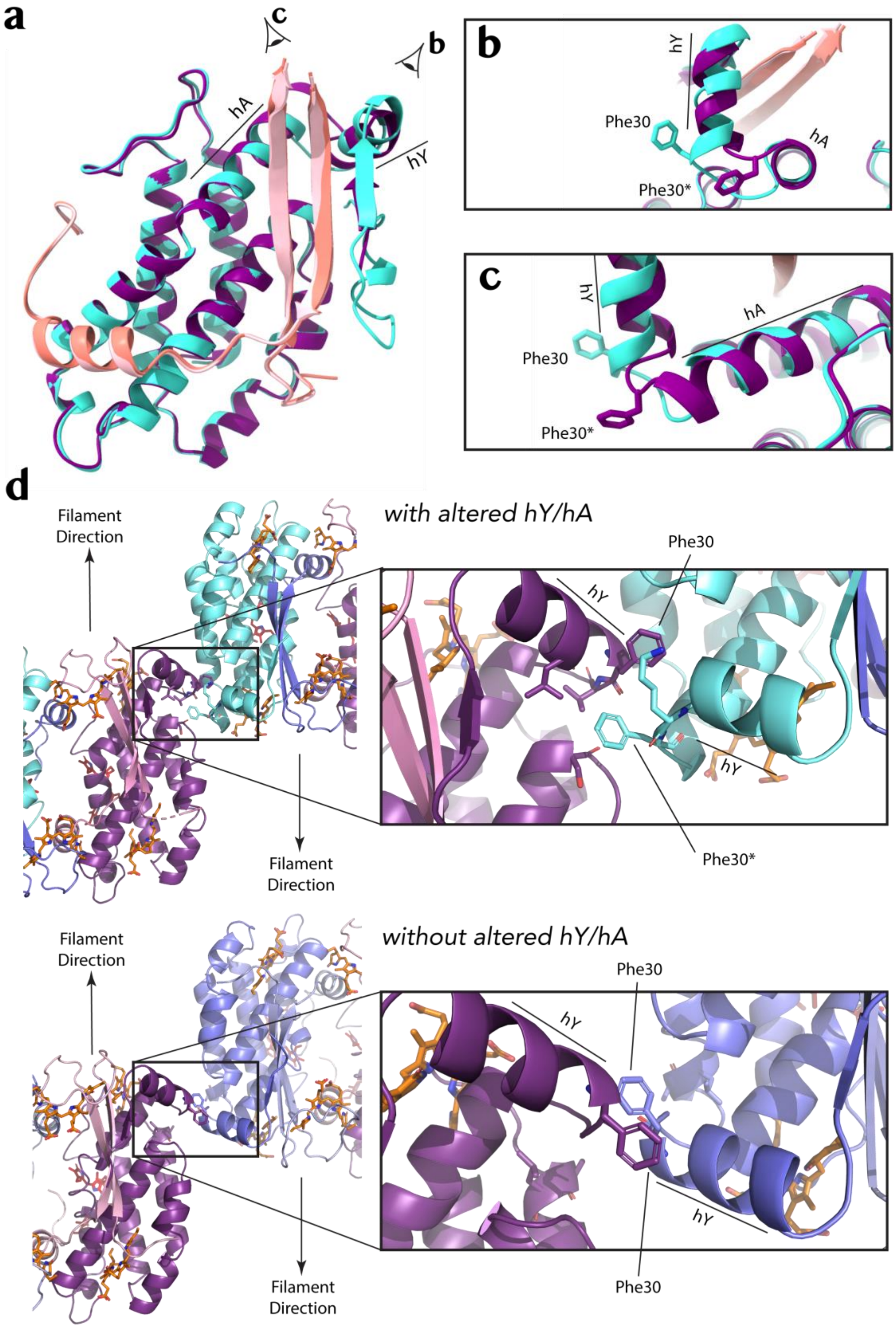


**Figure S3. Comparison of the crystal structures of *Ha*PE555A and *Ha*PE555B. a**. Superposition of the structure of *Ha*PE555A (α subunit salmon, β subunit cyan) and

*Ha*PE555B (α subunit light pink, β subunit purple). **b**. and **c**. show two enlargements of the superposition from the perspectives indicated by the eye symbols in **a**. These enlargements show the differences in α helices hY and hA with the associated change in the position of Phe30 in the two structures. **d**. The structural changes in helix hY and Phe30 alter the interfilament packing in *Ha*PE555B (top) compared with *Ha*PE555A (bottom), resulting in tighter packing in the former.

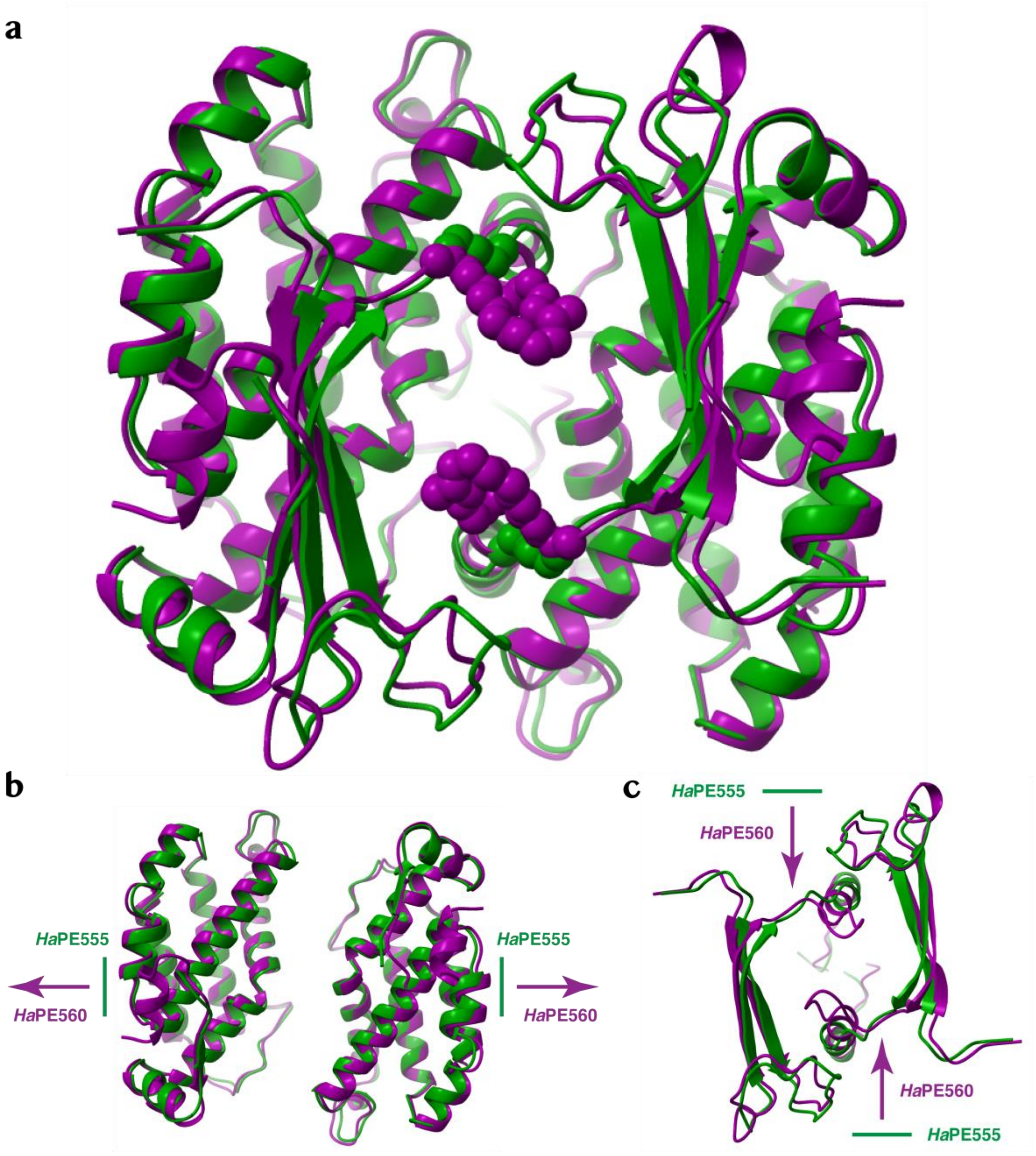


**Figure S4. Comparison of *open* and *open-braced* forms and the Poisson effect observed when comparing the *open-braced* form *Ha*PE560 with the *open* form *Ha*PE555. a**. Superposition of the *open-braced* form structure of *Ha*PE560 (purple) and the *open* form *Ha*PE555 (green). The *L1* loop, and the comparable path in *Ha*PE555, is displayed as a string of spheres to highlight how the incursion begins to fill the central solvent filled hole. The Poisson effect comprises a vertical compression coupled with a horizontal expansion of *Ha*PE560 with respect to *Ha*PE555. **b**. The horizontal displacements of the *Ha*PE560 β subunits resulting in a horizontal widening of the *open-braced* form compared to the *open*

form. **c**. the inward vertical compression of the two *Ha*PE560 α subunits when compared to *Ha*PE555 complete the Poisson effect.

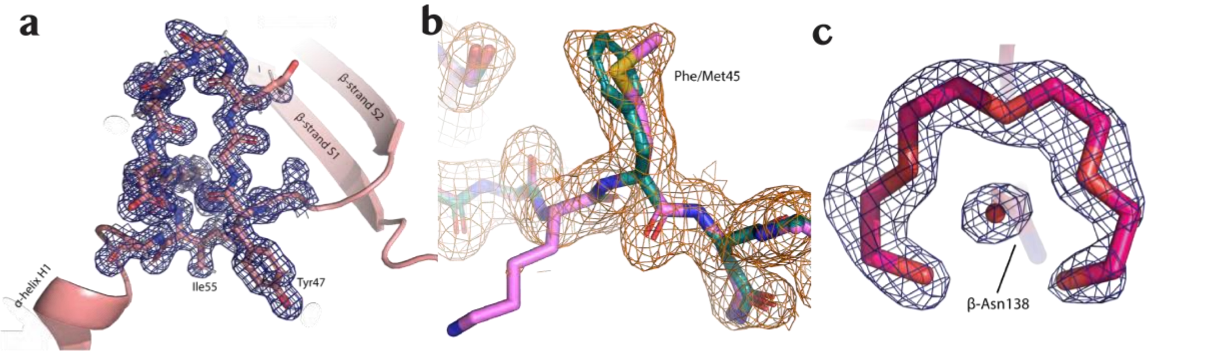


**Figure S5. Electron density for structural features of note.** **a**. Electron density map for the *Ha*PE560 α subunit *L1* loop overlayed on the structure. **b**. Electron density map for *Ha*PE555 showing the microheterogeneity at Met45/Phe45 in the α subunit. **c**. PEG molecule surrounds a water molecule that is hydrogen bonded to the side chain of Asn138 in the β subunit of *Ha*PE645.

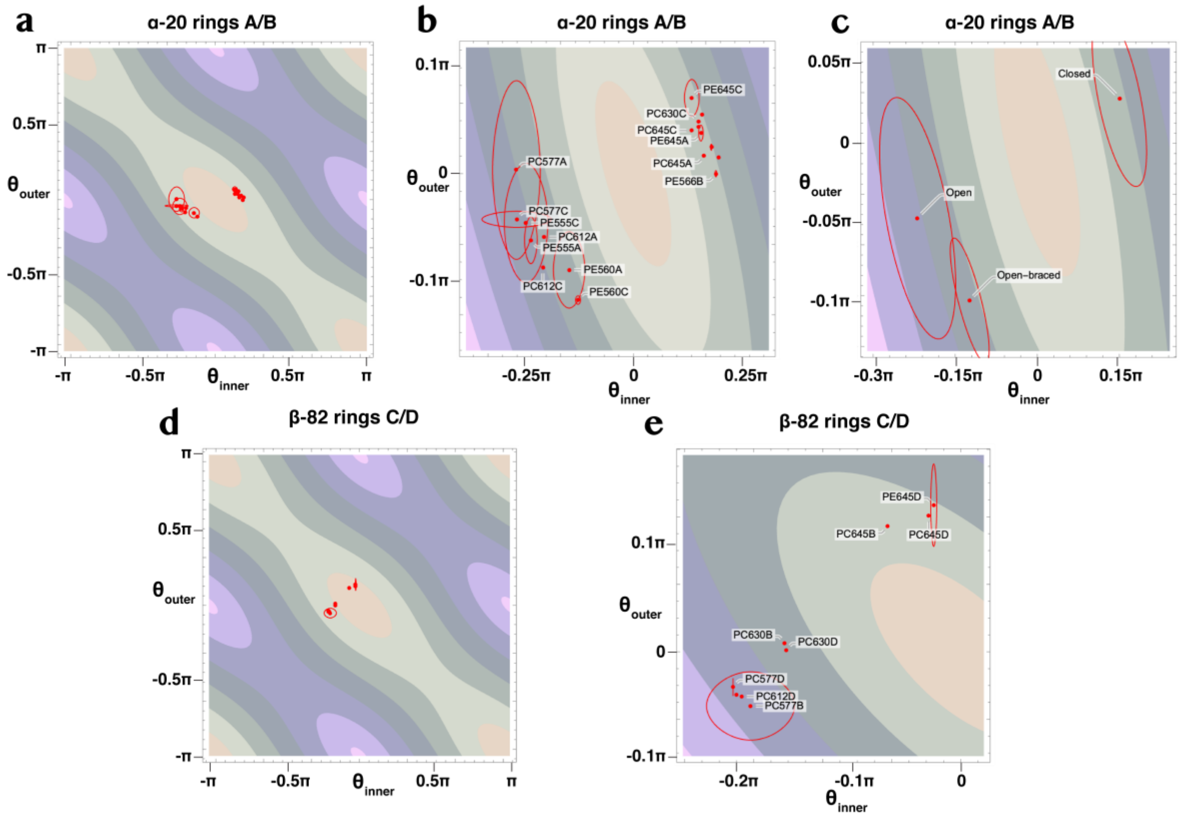


**Figure S6 Chromophore torsion angles.** **a-c**. The torsion angles between pyrrole rings A and B of the α chromophores. **a**. shows the full angular range where the *open* forms are clustered on the left of center while the *closed* forms are on the right. **b**. is a close up of the same plot as **a**. **c**. here the three quaternary structures: *open*, *open-braced* and *closed* separate into three distinct regions of dihedral space. **d-e**. The torsion angles between pyrrole rings C and D of the β82 chromophore. **d**. shows the full angular range while **e**. is a close up showing the orientations of ring D in the chromophore.

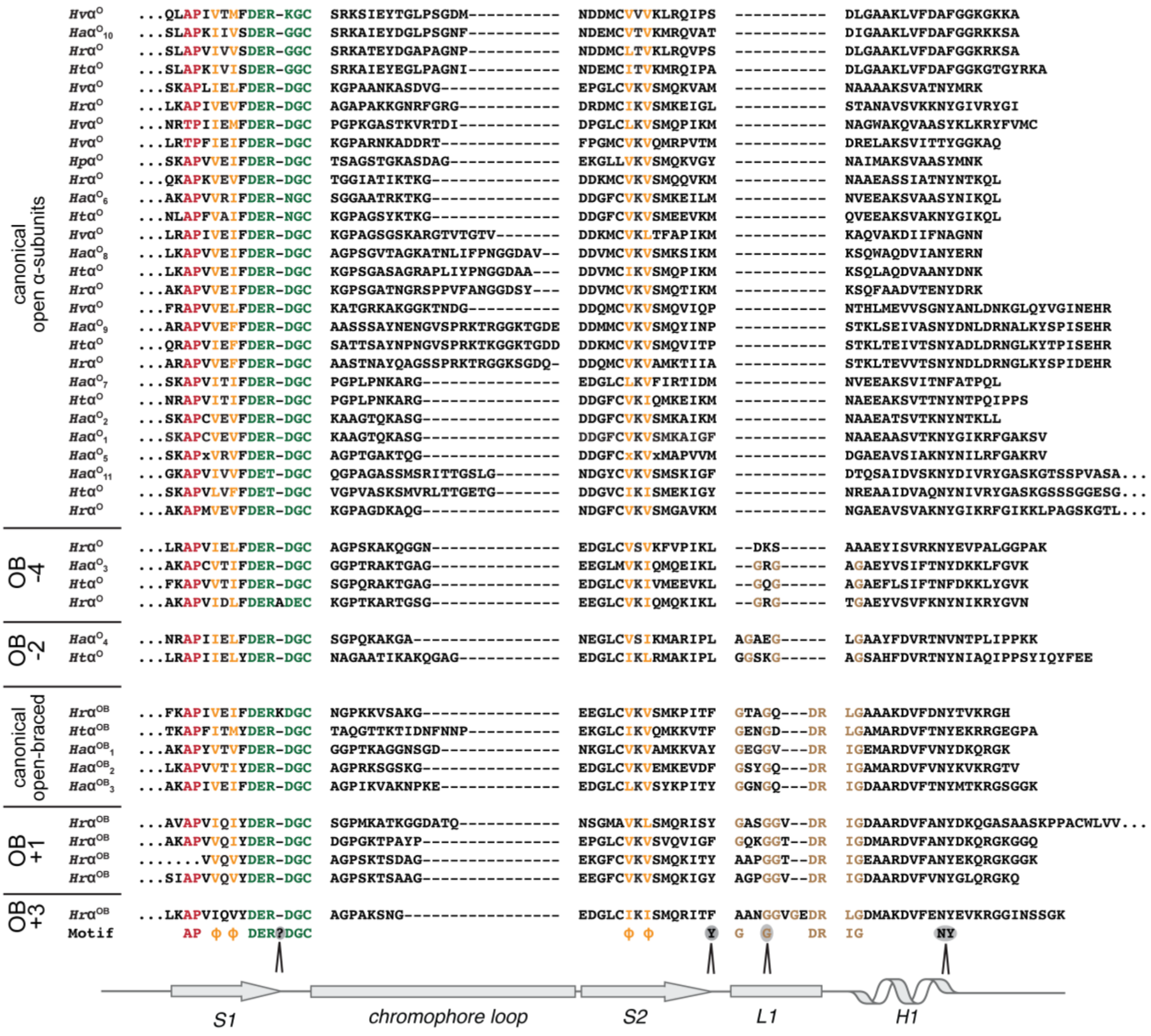


**Figure S7. Structure-based alignment of *open* form *Hemiselmis* α subunits highlighting *L1* insertions between β strand *S2* and the α helix.** An alignment of all *Hemiselmis* α subunits that show the *open* form characteristic (aspartic acid insertion two residues before the chromophore binding cysteine). The top block of sequences are canonical *open* form α subunits. The lower sequence blocks show sequences that contain an insertion between β strand *S2* and the α helix *H1* (see secondary structure at the bottom). These include the *open-braced* form as defined by the crystal structure of *Ha*PE560 (corresponding to sequence $Ha\alpha^{OB}{}_{1}$). The *L1* loop insertion for the *open-braced* form starts after an anchoring aromatic residue after β strand *S2* and terminates at an extra N-terminal turn on the α helix (RIG motif). The *L1 open-braced* motif is G-x-x-G-$x_{1\text{-}3}$-D-R. Ten sequences conform with this motif (clusters labelled canonical *open-braced*, OB +1 and OB +3). Six sequences exhibit a shorter insertion at this site (clusters labelled OB -2 and OB -4). Typically, these sequences have a G-$x_{1\text{-}2}$-G-x-G motif. The nature of the shorter insertion in these residues is currently unknown.

The first two letters of the sequence identifiers refer to the species: *Hv* - *H. virescens*; *Ha* – *H. andersenii*; *Hr* – *H. rufescens*; *Ht* – *H. tepida*; and *Hp* – *H. pacifica*. Color coding is as per Fig. 1a in the main manuscript.

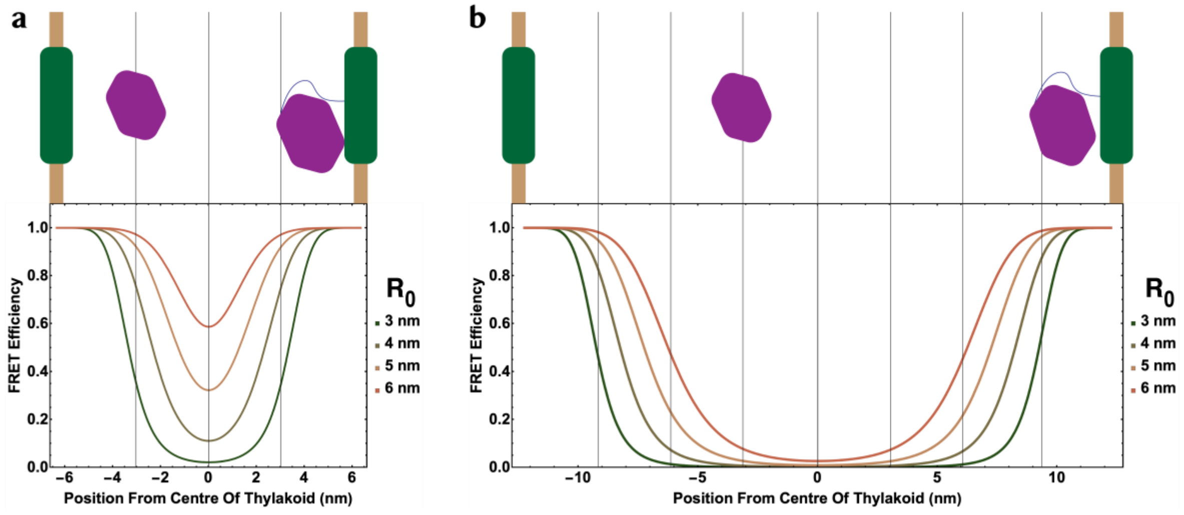


**Figure S8. Modelling the efficiency of energy transfer to the membrane systems as a function of the location of the adaptor, *Ha*PE645, within the thylakoid lumen.** **a**. Plot of FRET efficiency as a function of the position of *Ha*PE645 in a 12.7 nm wide thylakoid lumen. **b**. the same calculation for a 25 nm wide lumen, representing the half the extreme width of cryptophyte thylakoid compartments. Vertical lines represent a 'protein width'.

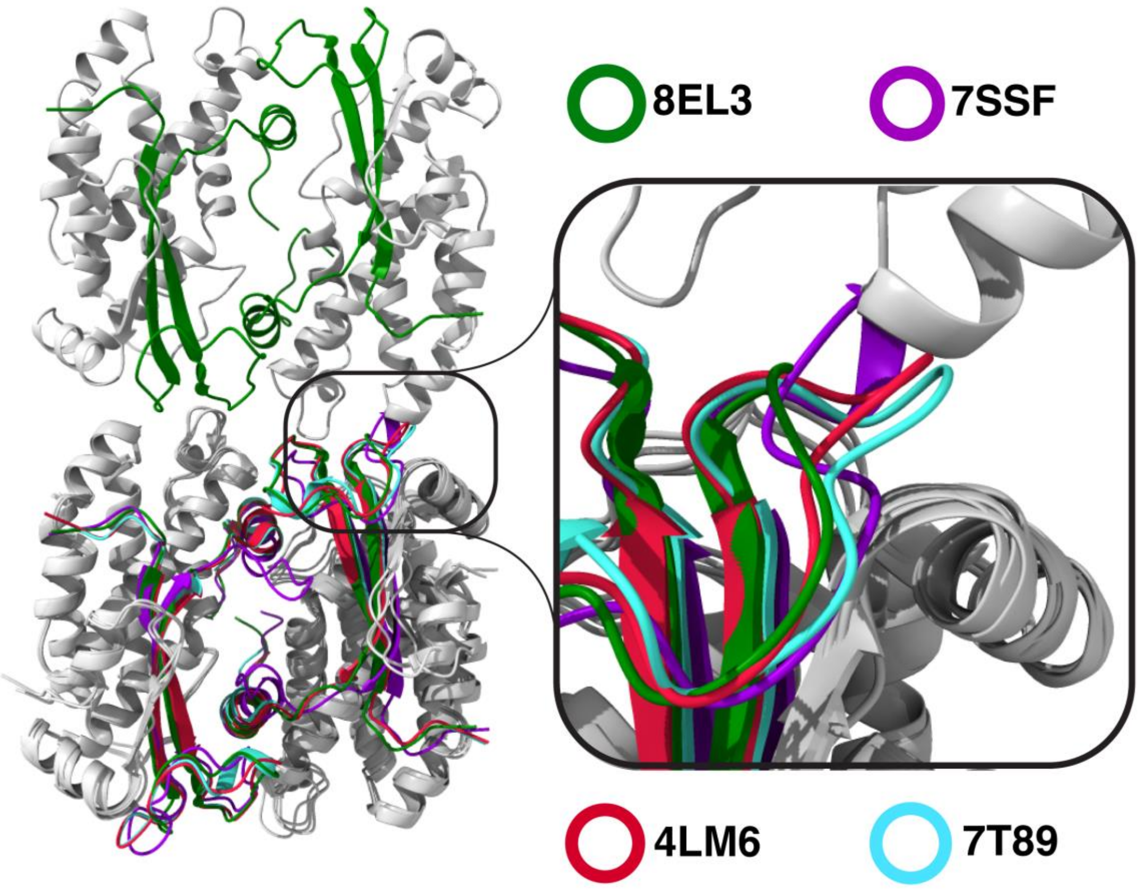


**Figure S9. Filament formation by *Ha*PE555 facilitated by α subunit chromophore loop structure.** Left panel shows the filament formed by *Ha*PE555 with a loose interface (8EL3). The upper molecule is *Ha*PE555 with the α subunits shown in green and the β subunits, gray. On the lower molecule, a complete PBP, $(\alpha\beta)_2$, has been superposed for each of: *Ha*PE560 (purple, 7SSF); *Hv*PC612 (red, 4LM6); and *Hp*PC577 (cyan, 7T89). As can be seen in the magnified view (right panel), all of these overlayed structures result in a steric clash with the neighboring molecule along the filament axis. The most severe clash is for *Ha*PE560, with smaller clashes for *Hv*PC612 and *Hp*PC577. Thus, without a structural rearrangement, only *Ha*PE555 is capable of forming the filament structures seen in all of its crystal forms. Chromophores are removed from view for clarity.

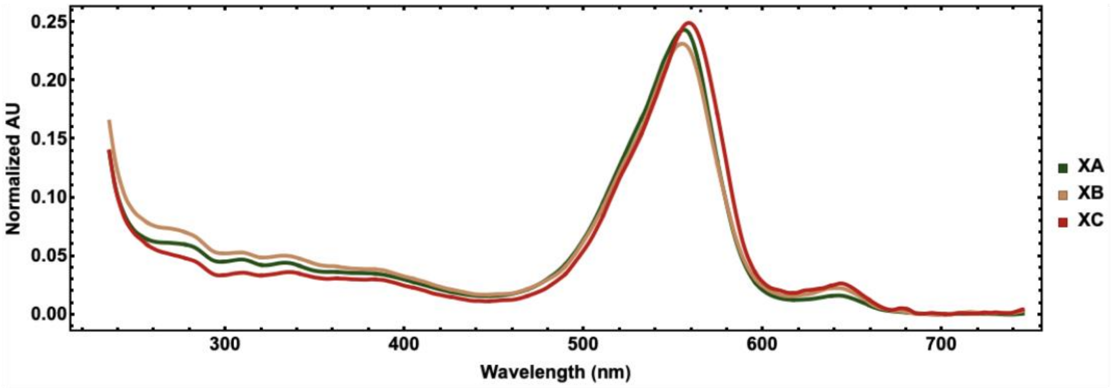


**Figure S10. Absorption spectra of unclassified peaks in chromatography.** Three minor peaks in the chromatogram in Fig. 1b (left) show absorption spectra that appear to indicate that they are mixtures of different spectrotypes (see Supplementary Text A).

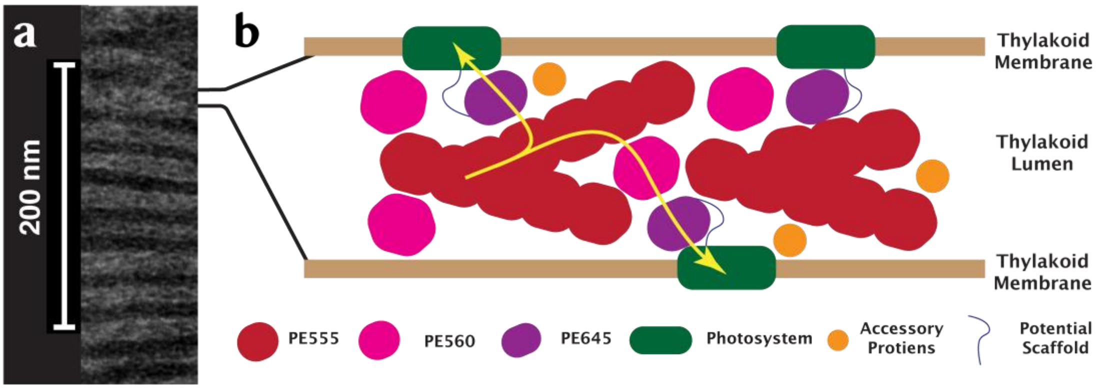


**Figure S11. Antenna model including *Ha*PE555 filaments.** **a**. electron micrograph showing electron dense material (dark bands) between the thylakoid membranes. **b**. model for the light harvesting antenna that includes filaments formed by *Ha*PE555.

**Table S1. Transcriptome α subunit sequences**

| α subunit | Strain | Accession Code |
|---|---|---|
| **HA$\alpha^{O}_{1}$** | — | *From PDB 4LMX* |
| **HA$\alpha^{O}_{2}$** | CCMP1180 | MMETSP1042-15997 |
| **HA$\alpha^{O}_{3}$** | CCMP44 | MMETSP1043-10158 |
| **HA$\alpha^{O}_{4}$** | CCMP1180 | MMETSP1042-7575 |
| **HA$\alpha^{O}_{5}$** | CCMP441 | MMETSP1043-6558 |
| **HA$\alpha^{O}_{6}$** | CCMP439 | MMETSP1041-8454 |
| **HA$\alpha^{O}_{7}$** | CCMP439 | MMETSP1041-10848 |
| **HA$\alpha^{O}_{8}$** | CCMP644 | MMETSP0043_2-9322 |
| **HA$\alpha^{O}_{9}$** | CCMP1180 | MMETSP1042-9535 |
| **HA$\alpha^{O}_{10}$** | CCMP1180 | MMETSP1042-7689 |
| **HA$\alpha^{O}_{11}$** | CCMP644 | MMETSP0043_2-10099 |
| **HA $\alpha^{OB}_{1}$** | CCMP1180 | MMETSP1042-7728 |
| **HA$\alpha^{OB}_{2}$** | CCMP441 | MMETSP1043-5984 |
| **HA$\alpha^{OB}_{3}$** | CCMP1180 | MMETSP1042-13440 |
| **HA$\alpha^{C}_{1}$** | CCMP1180 | MMETSP1042-4425 |
| **HA$\alpha^{C}_{2}$** | CCMP1180 | MMETSP1042-7955 |
| **HA$\alpha^{C}_{3}$** | CCMP1180 | MMETSP1042-22136 |
| **HA$\alpha^{C}_{4}$** | CCMP441 | MMETSP1043-8390 |
| **HA$\alpha^{C}_{5}$** | CCMP441 | MMETSP1043-11864 |
| **HA$\alpha^{C}_{6}$** | CCMP441 | MMETSP1043-6161 |
| **HA$\alpha^{C}_{7}$** | CCMP644 | MMETSP0043_2-17359 |
| **HA$\alpha^{C}_{8}$** | CCMP644 | MMETSP0043_2-29428 |

Common names of proteins and their accession codes.

**Table S2. Relative abundance of each light harvesting component by spectrotype**

| Gross Color Class | Percentage | Protein | % Of Gross Class | % Of Total |
|---|---|---|---|---|
| Pink | 85 ± 1% | *Ha*PE555A | 74% | 63 ± 1% |
| | | *Ha*PE560A | 16% | 14 ± 1% |
| | | Others | 10% | 8 ± 1% |
| Purple | 15 ± 1% | *Ha*PE645A | 53% | 8 ± 1% |
| | | Others | 47% | 7 ± 1% |

**Table S3. X-ray crystallographic data reduction and refinement**

| **Data collection**<br>***All (Outer Shell)*** | **8EL3**<br>***Ha*PE555** | **8EL4**<br>***Ha*PE555** | **8EL5**<br>***Ha*PE555** |
|---|---|---|---|
| Space group | P 1 21 1 | P 1 21 1 | P 1 21 1 |
| Cell dimensions | | | |
| *a*, *b*, *c* (Å) | 64.80, 76.84, 103.36 | 63.53, 71.26, 48.28 | 64.86, 75.60, 99.36 |
| α, β, γ (°) | 90, 110.73, 90 | 90, 108.47, 90 | 90, 110.30, 90 |
| Resolution (Å) | 1.57 (1.60 -1.57) | 1.73 (1.76-1.73) | 1.67 (1.70-1.67) |
| $R_{sym}$ or $R_{merge}$ | 0.095 (1.399) | 0.135 (2.797) | 0.091 (2.595) |
| $I / \sigma I$ | 10.5 (1.0) | 8.2 (0.7) | 9.5 (0.7) |
| Completeness (%) | 98.2 (81.9) | 100 (99.9) | 100 (99.7) |
| Redundancy | 6.4 (4.3) | 6.5 (6.7) | 6.7 (6.1) |
| $CC_{1/2}$ | 0.999 (0.40) | 0.997 (0.355) | 0.999 (0.406) |
| Wilson $B_{overall}$ (Å$^{-2}$) | 19.58 | 27.02 | 28.00 |
| **Refinement** | | | |
| Resolution (Å) | 1.57 | 1.73 | 1.67 |
| No. unique reflections | 129756 (5326) | 42700 (2325) | 104433 (5125) |
| $R_{work}$ / $R_{free}$ | 0.1701 / 0.2010 | 0.1877 / 0.2392 | 0.1858 / 0.2314 |
| No. protein chains | 8 | 4 | 8 |
| Non-protein/non-water components (not including alt. confs) | 4x AX9 12x PEB | 2x AX9 6x PEB | 3x GOL 4x AX9 12x PEB |
| No. atoms | | | |
| Protein | 17949 | 9386 | 19550 |
| Ligand/ion | 1596 | 798 | 1676 |
| Water | 695 | 192 | 340 |
| *B*-factors (Å$^{-2}$) | 23.34 | 35.85 | 39.11 |
| Protein | 23.21 | 36.19 | 39.55 |
| Ligand/ion | 20.60 | 32.60 | 35.27 |
| Water | 28.49 | 35.48 | 37.56 |
| **Geometry** | | | |
| Ramachandran Outliers | 0.00 | 0.00 | 0.00 |
| Ramachandran Favoured | 99.02 | 98.40 | 98.58 |
| Rotamer Outliers | 0.00 | 0.22 | 0.42 |
| C-beta Outliers | 0.00 | 0.00 | 0.00 |
| Clashscore | 2.74 | 3.77 | 4.91 |
| Rama Z-score | | | |
| whole | 0.03 | -1.02 | -0.29 |
| helix | -0.14 | -1.10 | -0.17 |
| sheet | 1.54 | 1.91 | 1.16 |
| loop | 0.26 | -0.31 | -0.21 |
| Molprobity Score | 0.90 | 1.17 | 1.18 |
| R.m.s. deviations | | | |
| Bond lengths (Å) | 0.013 | 0.013 | 0.013 |
| Bond angles (°) | 1.385 | 1.415 | 1.446 |

**Table S3 *Continued*. X-ray crystallographic data reduction and refinement**

| **Data collection** *All (Outer Shell)* | **8EL6** ***Ha*PE555** | **7SSF** ***Ha*PE560** | **7SUT** ***Ha*PE645** |
|---|---|---|---|
| Space group | P 1 21 1 | I 1 2 1 | P 1 21 1 |
| Cell dimensions | | | |
| *a*, *b*, *c* (Å) | 61.71, 70.00, 48.02 | 84.32, 67.97, 184.50 | 54.03, 80.98, 115.84 |
| α, β, γ (°) | 90, 110.31, 90 | 90, 99.33, 90 | 90, 92.21, 90 |
| Resolution (Å) | 1.83 (1.87-1.83) | 1.45 (1.47-1.45) | 1.49 (1.52-1.49) |
| $R_{sym}$ or $R_{merge}$ | 0.21 (1.539) | 0.095 (1.424) | 0.056 (1.117) |
| $I / \sigma I$ | 6.9 (1.2) | 12.5 (1.4) | 9.9 (0.9) |
| Completeness (%) | 100 (100) | 99.8 (99.3) | 93.4 (58.3) |
| Redundancy | 7.3 (7.3) | 7.4 (7.2) | 3.6 (2.2) |
| $CC_{1/2}$ | 0.994 (0.315) | 0.999 (0.546) | 0.998 (0.463) |
| Wilson $B_{overall}$ (Å$^{-2}$) | 18.02 | 14.93 | 19.89 |
| **Refinement** | | | |
| Resolution (Å) | 1.95 | 1.45 | 1.49 |
| No. unique reflections | 33901 (2072) | 181639 (8893) | 151716 (4645) |
| $R_{work}$ / $R_{free}$ | 0.1814 / 0.2554 | 0.1541 / 0.1921 | 0.1455 / 0.1747 |
| No. protein chains | 4 | 8 | 8 |
| Non-protein/non-water components (not including alt. confs) | 2x AX9 6x PEB | 4x AX9 12x PEB 3x TRS 1x CL | 4x DBV 4x AX9 2x PCB 6x PEB 2x PG4 1x BTB 1x CL |
| No. atoms | | | |
| Protein | 8437 | 15495 | 14867 |
| Ligand/ion | 798 | 1571 | 1443 |
| Water | 256 | 816 | 456 |
| *B*-factors (Å$^{-2}$) | 22.16 | 23.87 | 26.94 |
| Protein | 22.41 | 23.81 | 26.80 |
| Ligand/ion | 20.06 | 19.44 | 25.99 |
| Water | 21.57 | 28.93 | 30.69 |
| **Geometry** | | | |
| Ramachandran Outliers | 0.00 | 0.00 | 0.00 |
| Ramachandran Favoured | 97.70 | 98.54 | 98.07 |
| Rotamer Outliers | 0.67 | 0.48 | 0.12 |
| C-beta Outliers | 0.00 | 0.00 | 0.00 |
| Clashscore | 6.91 | 3.08 | 2.93 |
| Rama Z-score | | | |
| whole | -1.35 | -0.23 | -0.31 |
| helix | -1.23 | 0.14 | 0.04 |
| sheet | 0.53 | -0.05 | -0.01 |
| loop | -0.52 | -0.46 | -0.59 |
| Molprobity Score | 1.36 | 1.02 | 0.82 |
| R.m.s. deviations | | | |
| Bond lengths (Å) | 0.013 | 0.013 | 0.011 |
| Bond angles (°) | 1.504 | 1.415 | 1.317 |

**Table S4. Filament parameters for the four *Ha*PE555 crystal structures**

| PDB | ASU Content (# PBPs) | \|c\| / molecule (Å) | Condition (+25.25% PEG3350) |
|---|---|---|---|
| 8EL3 | 2 | 51.8 | 0.01M NaBr + 25.25% PEG3350 |
| 8EL5 | 2 | 49.7 | 0.2% (w/v) benzamidine HCl + 25.25% PEG3350 |
| 8EL4 | 1 | 48.3 | 0.01M sarcosine + 25.25% PEG3350 |
| 8EL6 | 1 | 48 | 23.6% PEG3350 |

The structures are ranked from that with the loosest interactions along the filament axis to that with the tightest interactions along the filament axis.

**Table S5. Chromophore complement for the three different spectrotypes**

| Protein | alpha | 50 | 82 | 158 |
|---|---|---|---|---|
| *Ha*PE555 | PEB | DBV | PEB | PEB |
| *Ha*PE560 | PEB | DBV | PEB | PEB |
| *Ha*PE645 | DBV | DBV | PCB/PEB | PEB |

**References and Notes:**

1. T. L. Richardson, The colorful world of cryptophyte phycobiliproteins. Journal of Plankton Research **44**, 806-818 (2022).

2. J. M. Archibald, Cryptomonads. Curr Biol **30**, R1114-R1116 (2020).

3. H. W. Rathbone, K. A. Michie, M. J. Landsberg, B. R. Green, P. M. G. Curmi, Scaffolding proteins guide the evolution of algal light harvesting antennas. Nat Commun **12**, 1890 (2021).

4. X. You et al., In situ structure of the red algal phycobilisome-PSII-PSI-LHC megacomplex. Nature **616**, 199-206 (2023).

5. N. Adir, S. Bar-Zvi, D. Harris, The amazing phycobilisome. Biochim Biophys Acta Bioenerg **1861**, 148047 (2020).

6. K. E. Apt, J. L. Collier, A. R. Grossman, Evolution of the phycobiliproteins. J Mol Biol **248**, 79-96 (1995).

7. A. B. Doust et al., Developing a structure-function model for the cryptophyte phycoerythrin 545 using ultrahigh resolution crystallography and ultrafast laser spectroscopy. J Mol Biol **344**, 135-153 (2004).

8. K. E. Wilk et al., Evolution of a light-harvesting protein by addition of new subunits and rearrangement of conserved elements: crystal structure of a cryptophyte phycoerythrin at 1.63-A resolution. Proc Natl Acad Sci U S A **96**, 8901-8906 (1999).

9. S. J. Harrop et al., Single-residue insertion switches the quaternary structure and exciton states of cryptophyte light-harvesting proteins. Proc Natl Acad Sci U S A **111**, E2666-2675 (2014).

10. K. A. Michie et al., Molecular structures reveal the origin of spectral variation in cryptophyte light harvesting antenna proteins. Protein Sci **32**, e4586 (2023).

11. C. O'hEocha, M. Raftery, Phycoerythrins and phycocyanins of cryptomonads. Nature **184**, 1049-1051 (1959).

12. F. T. Haxo, D. C. Fork, Photosynthetically active accessory pigments of cryptomonads. Nature **184**, 1051-1052 (1959).

13. M. B. Allen, E. C. Dougherty, L. J. Mc, Chromoprotein pigments of some cryptomonad flagellates. Nature **184**, 1047-1049 (1959).

14. A. N. Glazer, G. J. Wedemayer, Cryptomonad biliproteins - An evolutionary perspective. Photosynthesis Research **46**, 93-105 (1995).

15. E. Gantt, in Biochemistry and Physiology of Protozoa (Second Edition), M. Levandowsky, S. H. Hutner, Eds. (Academic Press, 1979), pp. 121-137.

16. D. R. A. Hill, K. S. Rowan, The biliproteins of the Cryptophyceae. Phycologia **28**, 455-463 (1989).

17. M. J. Broughton, C. J. Howe, R. G. Hiller, Distinctive organization of genes for light-harvesting proteins in the cryptophyte alga Rhodomonas. Gene **369**, 72-79 (2006).

18. J. Jenkins, R. G. Hiller, J. Speirs, J. Godovac-Zimmermann, A genomic clone encoding a cryptophyte phycoerythrin alpha-subunit. Evidence for three alpha-subunits and an N-terminal membrane transit sequence. FEBS Lett **273**, 191-194 (1990).

19. E. Morschel, W. Wehrmeyer, Multiple forms of phycoerythrin-545 from Cryptomonas maculata. Arch Microbiol **113**, 83-89 (1977).

20. E. Morschel, W. Wehrmeyer, Cryptomonad biliprotein: phycocyanin-645 from a Chroomonas species. Arch Microbiol **105**, 153-158 (1975).

21. A. N. Glazer, G. Cohen-Bazire, A comparison of cryptophytan phycocyanins. Arch Microbiol **104**, 29-32 (1975).

22. C. Brooks, E. Gantt, Comparison of phycoerythrins (542, 566nm) from cryptophycean algae. Arch Mikrobiol **88**, 193-204 (1973).

23. B. A. Curtis et al., Algal genomes reveal evolutionary mosaicism and the fate of nucleomorphs. Nature **492**, 59-65 (2012).

24. T. Kieselbach, O. Cheregi, B. R. Green, C. Funk, Proteomic analysis of the phycobiliprotein antenna of the cryptophyte alga Guillardia theta cultured under different light intensities. Photosynth Res **135**, 149-163 (2018).

25. L. C. Spangler, M. Yu, P. D. Jeffrey, G. D. Scholes, Controllable Phycobilin Modification: An Alternative Photoacclimation Response in Cryptophyte Algae. ACS Cent Sci **8**, 340-350 (2022).

26. H. Scheer, K. H. Zhao, Biliprotein maturation: the chromophore attachment. Mol Microbiol **68**, 263-276 (2008).

27. K. E. Overkamp et al., Insights into the biosynthesis and assembly of cryptophycean phycobiliproteins. J Biol Chem **289**, 26691-26707 (2014).

28. L. K. Anderson, C. M. Toole, A model for early events in the assembly pathway of cyanobacterial phycobilisomes. Mol Microbiol **30**, 467-474 (1998).

29. C. Lichtle, Effects of Nitrogen Deficiency and Light of High-Intensity on Cryptomonas-Rufescens (Cryptophyceae) .1. Cell and Photosynthetic Apparatus Transformations and Encystment. Protoplasma **101**, 283-299 (1979).

30. M. A. Faust, E. Gantt, Effect of Light-Intensity and Glycerol on Growth, Pigment Composition, and Ultrastructure of Chroomonas Sp. Journal of Phycology **9**, 489-495 (1973).

31. E. Gantt, M. R. Edwards, Provasol.L, Chloroplast Structure of Cryptophyceae - Evidence for Phycobiliproteins within Intrathylakoidal Spaces. Journal of Cell Biology **48**, 280-& (1971).

32. T. Mirkovic, K. E. Wilk, P. M. Curmi, G. D. Scholes, Phycobiliprotein diffusion in chloroplasts of cryptophyte Rhodomonas CS24. Photosynth Res **100**, 7-17 (2009).

33. R. Kana, O. Prasil, C. W. Mullineaux, Immobility of phycobilins in the thylakoid lumen of a cryptophyte suggests that protein diffusion in the lumen is very restricted. FEBS Lett **583**, 670-674 (2009).

34. K. M. Heidenreich, T. L. Richardson, Photopigment, Absorption, and Growth Responses of Marine Cryptophytes to Varying Spectral Irradiance. J Phycol **56**, 507-520 (2020).

35. C. D. van der Weij-De Wit et al., Phycocyanin Sensitizes both Photosystem I and Photosystem II in Cryptophyte Chroomonas CCMP270 Cells. Biophysical Journal **94**, 2423-2433 (2008).

36. L. S. Zhao et al., Structural basis and evolution of the photosystem I-light-harvesting supercomplex of cryptophyte algae. Plant Cell **35**, 2449-2463 (2023).

37. C. D. Martin, R. G. Hiller, Subunits and chromophores of a type I phycoerythrin from a Chroomonas sp. (Cryptophyceae). Biochimica et Biophysica Acta (BBA) - General Subjects **923**, 88-97 (1987).

38. R. G. Hiller, C. D. Martin, Multiple forms of a type I phycoerythrin from a Chroomonas sp. (Cryptophyceae) varying in subunit composition. Biochim Biophys Acta **923**, 98-102 (1987).

39. N. P. Cowieson et al., MX1: a bending-magnet crystallography beamline serving both chemical and macromolecular crystallography communities at the Australian Synchrotron. J Synchrotron Radiat **22**, 187-190 (2015).

40. D. Aragao et al., MX2: a high-flux undulator microfocus beamline serving both the chemical and macromolecular crystallography communities at the Australian Synchrotron. J Synchrotron Radiat **25**, 885-891 (2018).

41. K. Katoh, K. Misawa, K. Kuma, T. Miyata, MAFFT: a novel method for rapid multiple sequence alignment based on fast Fourier transform. Nucleic Acids Res **30**, 3059-3066 (2002).

42. J. J. Almagro Armenteros et al., Detecting sequence signals in targeting peptides using deep learning. Life Sci Alliance **2**, e201900429 (2019).

43. D. N. Perkins, D. J. Pappin, D. M. Creasy, J. S. Cottrell, Probability-based protein identification by searching sequence databases using mass spectrometry data. Electrophoresis **20**, 3551-3567 (1999).

44. C. C. P. N. 4, The CCP4 suite: programs for protein crystallography. Acta Cryst. D **50**, 760-763 (1994).

45. P. D. Adams et al., PHENIX: building new software for automated crystallographic structure determination. Acta Crystallogr D Biol Crystallogr **58**, 1948-1954 (2002).

46. P. Emsley, K. Cowtan, Coot: model-building tools for molecular graphics. Acta Crystallogr D Biol Crystallogr **60**, 2126-2132 (2004).

47. W. L. DeLano, The PyMOL User's Manual. (DeLano Scientific, San Carlos, CA, USA, 2002).

48. T. D. Goddard et al., UCSF ChimeraX: Meeting modern challenges in visualization and analysis. Protein Sci **27**, 14-25 (2018).

49. C. C. Jumper, I. H. M. van Stokkum, T. Mirkovic, G. D. Scholes, Vibronic Wavepackets and Energy Transfer in Cryptophyte Light-Harvesting Complexes. J Phys Chem B **122**, 6328-6340 (2018).

50. K. E. Overkamp et al., Chromophore composition of the phycobiliprotein Cr-PC577 from the cryptophyte Hemiselmis pacifica. Photosynth Res **122**, 293-304 (2014).

51. A. J. Laos et al., Cooperative Subunit Refolding of a Light-Harvesting Protein through a Self-Chaperone Mechanism. Angew Chem Int Ed Engl **56**, 8384-8388 (2017).

52. V. May, O. Kühn, Charge and Energy Transfer Dynamics in Molecular Systems. (Wiley, 2011).

53. P. M. Krasilnikov, D. V. Zlenko, I. N. Stadnichuk, Rates and pathways of energy migration from the phycobilisome to the photosystem II and to the orange carotenoid protein in cyanobacteria. FEBS Lett **594**, 1145-1154 (2020).

54. J. Grabowski, E. Gantt, PHOTOPHYSICAL PROPERTIES OF PHYCOBILIPROTEINS FROM PHYCOBILISOMES: FLUORESCENCE LIFETIMES, QUANTUM YIELDS, AND POLARIZATION SPECTRA. Photochemistry and Photobiology **28**, 39-45 (1978).

55. H. W. Rathbone, J. A. Davis, P. M. Curmi, in Photosynthesis in Algae, A. W. Larkum, J. A. Raven, A. Grossman, Eds. (Springer Verlag, 2020).

56. R. MacColl, D. S. Berns, O. Gibbons, Characterization cryptomonad phycoerythrin and phycocyanin. Arch Biochem Biophys **177**, 265-275 (1976).

57. A. N. Glazer, G. Cohen-Bazire, R. Y. Stanier, Characterization of phycoerythrin from a Cryptomonas sp. Arch Mikrobiol **80**, 1-18 (1971).

58. R. MacColl, W. Habig, D. S. Berns, Characterization of phycocyanin from Chromonas species. J Biol Chem **248**, 7080-7086 (1973).

59. C. J. Grisdale, D. R. Smith, J. M. Archibald, Relative Mutation Rates in Nucleomorph-Bearing Algae. Genome Biol Evol **11**, 1045-1053 (2019).
